# Oligodendrocytes and neurons contribute to amyloid-β deposition in Alzheimer’s disease

**DOI:** 10.1101/2023.12.11.570514

**Authors:** Andrew Octavian Sasmita, Constanze Depp, Taisiia Nazarenko, Ting Sun, Sophie B. Siems, Xuan Yu, Carolin Boehler, Erinne Cherisse Ong, Bastian Bues, Lisa Evangelista, Barbara Morgado, Zoe Wu, Torben Ruhwedel, Swati Subramanian, Friederike Börensen, Katharina Overhoff, Lena Spieth, Stefan A. Berghoff, Katherine Rose Sadleir, Robert Vassar, Simone Eggert, Sandra Goebbels, Takashi Saito, Takaomi Saido, Wiebke Möbius, Gonçalo Castelo-Branco, Hans-Wolfgang Klafki, Oliver Wirths, Jens Wiltfang, Sarah Jäkel, Riqiang Yan, Klaus-Armin Nave

**Affiliations:** Department of Neurogenetics, Max Planck Institute for Multidisciplinary Sciences, Göttingen, Germany; International Max Planck Research School for Neurosciences, Göttingen, Germany; Laboratory of Molecular Neurobiology, Department of Biochemistry and Biophysics, Karolinska Institutet, Stockholm, Sweden; School of Biochemistry and Cell Biology, Biosciences Institute, University College Cork, Cork, Ireland; Institute for Stroke and Dementia Research, Klinikum Der Universität München, Ludwig-Maximilians-Universität, Munich, Germany; Department of Psychiatry and Psychotherapy, University Medical Center, Georg-August University, Göttingen, Germany; Electron Microscopy Core Unit, Max Planck Institute Multidisciplinary Sciences, Göttingen, Germany; Ken and Ruth Davee Department of Neurology, Northwestern University Feinberg School of Medicine, Chicago, USA; Mesulam Center for Cognitive Neurology and Alzheimer’s Disease, Northwestern University Feinberg School of Medicine, Northwestern University, Chicago, USA; Laboratory for Proteolytic Neuroscience, RIKEN Center for Brain Science, Wako, Saitama, Japan; German Center for Neurodegenerative Diseases (DZNE), Göttingen, Germany; Munich Cluster for System Neurology (SyNergy) Munich; Department of Neuroscience, UConn Health, Farmington, USA

**Author notes:** **Correspondence** Communication and request for materials, data, or code pertaining to this study should be addressed to, or. These authors contributed equally (Andrew Octavian Sasmita, Constanze Depp).

## Abstract

In Alzheimer’s disease (AD), amyloid-β (Aβ) is thought to be of neuronal origin. However, in single-cell RNAseq datasets from mouse and human, we found transcripts of amyloid precursor protein (APP) and the amyloidogenic-processing machinery equally abundant in oligodendrocytes (OLs). By cell-type-specific deletion of Bace1 in a humanized knock-in AD model, *APP*^*NLGF*^, we demonstrate that almost a third of cortical Aβ deposited in plaques is derived from OLs. However, excitatory projection neurons must provide a threshold level of Aβ production for plaque deposition to occur and for oligodendroglial Aβ to co-aggregate. Indeed, very few plaques are deposited in the absence of neuronally-derived Aβ, although soluble Aβ species are readily detected, especially in subcortical white matter. Our data identify OLs as a source of Aβ in vivo and further underscore a non-linear relationship between cellular Aβ production and resulting plaque formation. Ultimately, our observations are relevant for therapeutic strategies aimed at disease prevention in AD.

## Main text

In Alzheimer’s disease (AD) and its mouse models, amyloid-β (Aβ) production has primarily been attributed to excitatory neurons (ExNs)^1^, despite emerging evidence that other cell types, such as inhibitory interneurons or glial cells, might contribute to Aβ production^2,3^. Cultured oligodendrocytes (OLs) are capable of generating detectable levels of Aβ in *vitro*^4-6^. Since OL lineage cells are abundantly present in both gray and white matter (WM) and myelin alterations have been implicated in AD ^7-10^, we asked whether OLs would directly contribute to Aβ plaque burden *in vivo*.

### Amyloidogenic genes are amply expressed by OL-lineage cells

We first interrogated multiple *in vivo* single cell RNA-Seq (scRNA-Seq) and single nuclear RNA-Seq (snRNA-Seq) datasets of wild-type (WT) mouse^10-12^ and healthy control human^13-15^ nervous tissue for expression of amyloidogenic pathway genes (*APP, BACE1, PSEN1, PSEN2*) in all major brain cell types, including OLs (**Fig. 1, Extended Data Fig. 1a-b**). Depending on the sequencing technology and tissue input, positive cell rates of amyloidogenic transcripts varied but expression levels were comparable between excitatory neurons (ExNs) and OLs, both mature (MOLs) and newly-formed (NFOLs) (**Extended Data Fig. 1c-d**). We further validated the expression of APP and BACE1 in murine OLs within their cell soma and processes *in vitro* (**Extended Data Fig. 2a**). By *in situ* hybridization (ISH) in human cortical tissue, we found that around 50% of all gray matter OLs express significant levels of *APP* mRNA in both AD cases and controls (**Extended Data Fig. 2b-c**). Thus, both mouse and human OLs express the essential genes for Aβ generation *in vivo*.

**Figure 1.**
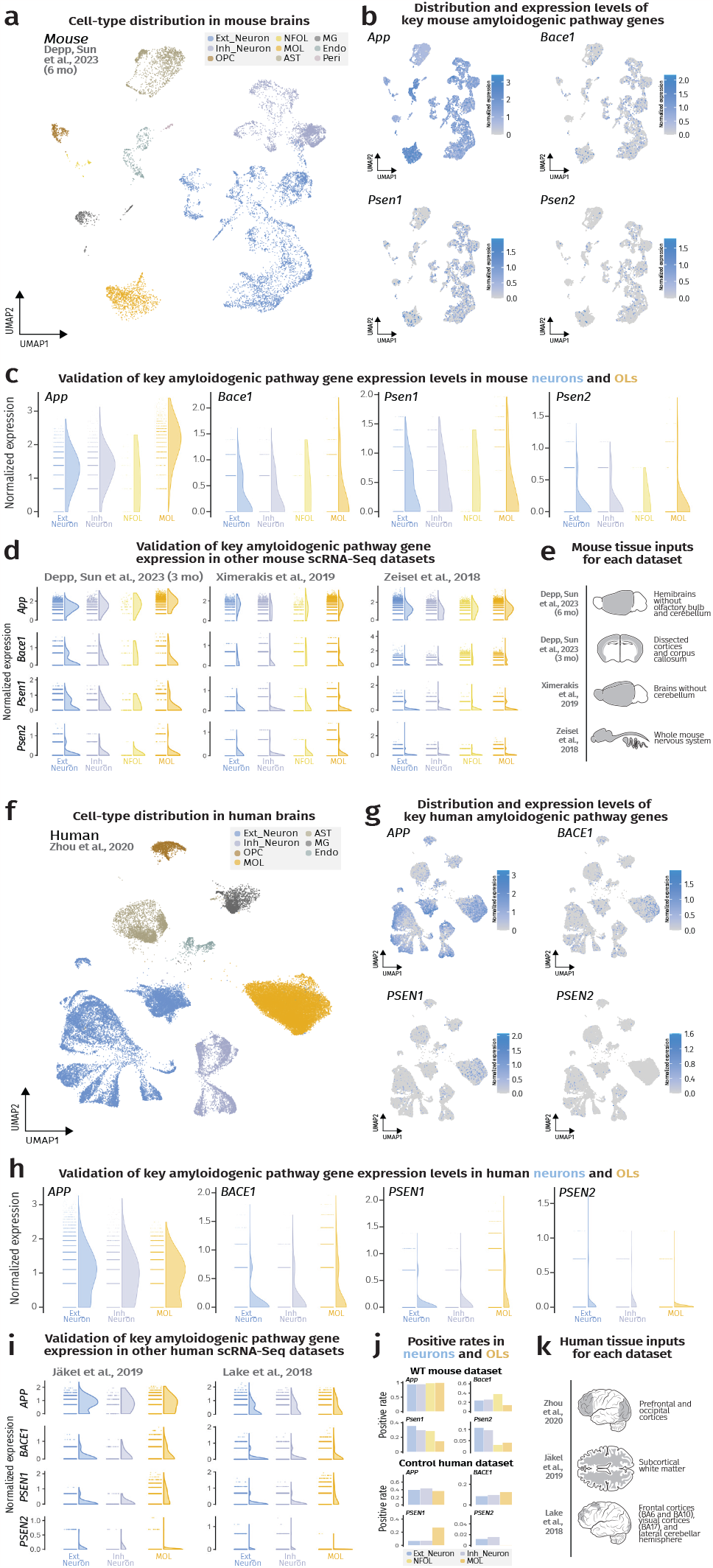
OLs abundantly express key amyloidogenic pathway genes as assessed by scRNA-Seq and snRNA-Seq. (**a**) UMAP visualization of cell types from a 6-month-old mouse brain snRNA-Seq dataset^10^. (**b**) Feature plots showcasing expression of key amyloidogenic genes (*App, Bace1, Psen1*, and *Psen2*) across all cell types in WT mouse brains. (**c**) Expression level half violin plots of key amyloidogenic genes in neurons and OLs of mouse brains normalized by the SCTransform method, highlighting the comparable expression of all genes between neurons and OLs. (**d**) Expression level half violin plots of key amyloidogenic genes in neurons and OLs with SCTransform normalization method from additional mouse datasets^10-12^. (**e**) Mouse nervous tissue inputs for sequencing from each study shown. (**f**) UMAP visualization of cell types from a human brain snRNA-Seq dataset^13^. (**g**) Feature plots showcasing expression of key amyloidogenic genes (*APP, BACE1, PSEN1*, and *PSEN2*) across all cell types in control human brains. (**h**) Expression level half violin plots of key amyloidogenic genes in neurons and OLs of human brains with SCTransform normalization method, highlighting the comparable expression of all genes between neurons and OLs. (**i**) Expression level half violin plots of key amyloidogenic genes in neurons and OLs normalized by SCTransform normalization method from additional human datasets^14,15^. (**j**) Positive rate barplots of APP processing genes in mouse and human nervous tissue inputs. (**k**) Human nervous tissue inputs for sequencing from each study shown. (**c, d, h, i**) Half violins represent aggregated expression levels of respective genes from each cell type and data points refer to individual expression levels from single cells or nuclei normalized by SCTransform. The results published here are based on data obtained from GEO and the AD Knowledge Portal.

### *Bace1* cKO resulted in an OL- and ExN-specific *Bace1* ablation and modulated APP processing

Next, we created novel AD mouse lines to assess Aβ contribution from OLs and ExNs separately (**Fig. 2a**). For this, we employed *APP*^*NLGF*^ knock-in mice that express a humanized and triple-mutated amyloid precursor protein (*APP*) in the endogenous *App* locus^16^ to circumvent potential transgenic mouse artifacts. These mice were crossed with recently generated *Bace1*^*fl/fl*^ mice to conditionally knock-out *Bace1* (*Bace1* cKO)^17^, the rate-limiting enzyme in Aβ generation, using cell-type-specific *Cre* drivers. The ability to process Aβ from APP was abrogated specifically in OLs using the *Cnp-Cre* driver line^18^ or ExNs using the *Nex-Cre* driver line^19^. We termed the resultant triple mutant mice *OL-Bace1*^*cKO*^*;AD* and *ExN-Bace1*^*cKO*^*;AD* respectively and compared them to non-*Cre* controls termed *Control;AD*.

**Figure 2.**
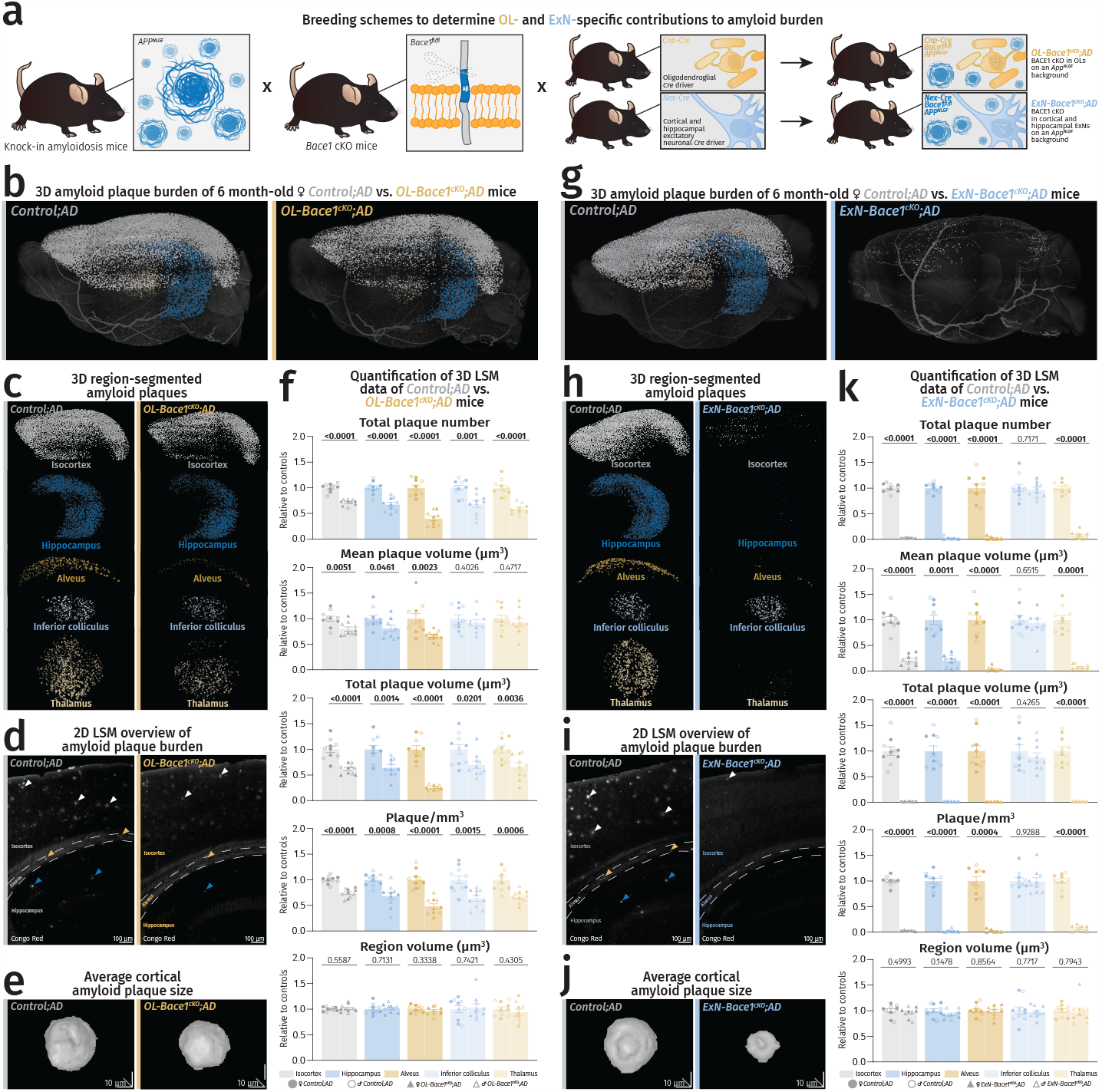
OLs contribute to Aβ burden primarily derived from ExNs *in vivo*. (**a**) Mouse breeding setup to investigate the OL- and ExN-specific contributions to Aβ burden. (**b-f**) Light sheet microscopy data of plaque burden (Congo red) comparing 6-month-old *OL-Bace1*^*cKO*^*;AD* mice to age- and sex-matched littermate controls. (**g-k**) Light sheet microscopy data of plaque burden (Congo red) comparing 6-month-old *ExN-Bace1*^*cKO*^*;AD* mice to age- and sex-matched littermate controls. (**b-k**) Color-region allocation is as follows: White-isocortex, blue-hippocampus, yellow-alveus, pastel blue-inferior colliculus, pastel yellow-thalamus. (**b, g**) LSM 3D visualization of control and cKO hemibrains. (**c, h**) Brain region-segmented plaques of control and cKO hemibrains. (**d, i**) LSM 2D single plane of control and cKO hemibrains. Arrows point to plaques with colors indicating specific regions. (**e, j**) LSM 3D renders of representative cortical Aβ plaques of control and cKO hemibrains. (**f, k**) Quantification of LSM data between controls (n=5/sex) and cKOs (n=5/sex). Normalization of cKO data points was performed to sex-matched controls. Circles represent controls and triangles represent cKOs. Filled shapes represent male and hollowed shapes represent female mice. For each parameter, unpaired, two-tailed Student’s t-test was performed (p-values indicated in graphs with significance highlighted in bold) comparing cKOs to controls. Bars represent means with SEM and individual data points displayed. Raw data is available in **Extended Data Table 1**.

We assessed *Cnp-Cre* specificity using a stop-flox tdTomato reporter mouse line^20^ as transient *Cnp-Cre* activity has been detected with sensitive reporter functions in the neuronal lineage to varying degrees^21,22^. In concordance with recent findings^23^, only a very low percentage of cortical (0.756% ± 0.057) and hippocampal (0.468% ± 0.111) neurons were tdTomato-positive (**Extended Data Fig. 3**). We then validated the cell-type-specific *Bace1* transcript reduction using ISH, whereby *Bace1* transcripts were massively reduced in the intended target-cell type (OLs for *Cnp-Cre* and ExN for *Nex-Cre* expression) (**Extended Data Fig. 4**). Importantly, we confirmed that ExN *Bace1* transcript levels were unaffected in *OL-Bace1*^*cKO*^*;AD* animals (**Extended Data Fig. 4a-d**, **h-m**).

We turned to Western blot analysis to validate APP processing alterations associated with *Bace1* cKO (**Extended Data Fig. 5**). Full-length APP (FL-APP) levels were 40% lower in control *APP*^*NLGF*^ lysates compared to WT brains, most likely because mutated APP preferably underwent β-cleavage to produce β C-terminal fragments (β-CTFs). As expected, both *Bace1* cKOs in ExNs and OLs resulted in a region-dependent depletion of BACE1, reflecting local differences in the neuron to OL ratio. Of note, WM tracts harbor a substantial amount of axoplasm containing neuronally-expressed BACE1^24^, explaining the reduction of this protein in the WM fraction from *ExN-Bace1*^*cKO*^*;AD* mice. Confirming the successful targeting of BACE1, cell-type-specific losses of BACE1 diminished β-CTFs in the cKOs and restored FL-APP to nearly baseline WT amounts. Levels of presenilin-1, one of the catalytic subunits of gamma secretase, remained unchanged in the absence of BACE1.

### OLs contribute to Aβ burden *in vivo* with strongest effects observed in the white matter

We then utilized light sheet microscopy (LSM) for *in toto* imaging of Congo red-stained amyloid plaques in *OL-Bace1*^*cKO*^*;AD* and *ExN-Bace1*^*cKO*^*;AD* mouse hemibrains at 6 months in both sexes as sexual dimorphism in this model was evident (**Extended Data Fig. 6**). Analyzed hemibrains were then segmented into regions of interest (ROIs), including the cortex and hippocampus, the alveus as a representative WM tract, and lastly the thalamus and inferior colliculus as regions that do not show *Nex-Cre* recombination^19^.

*OL-Bace1*^*cKO*^*;AD* mice exhibited a reduced plaque burden indicating that OLs actively contribute to amyloid burden *in vivo*. (**Fig. 2b-f**). At 6 months of age, *OL-Bace1*^*cKO*^*;AD* mice accumulated approximately 30% less plaques when compared to respective controls in both sexes. The decrease in plaque amount and, more importantly, in plaque size was greatest in the alveus, a WM tract that develops amyloid pathology. In addition, microgliosis was proportional to Aβ plaque pathology (**Extended Data Fig. 7**).

### Deletion of *Bace1* in cortical and hippocampal ExNs reduced cerebral Aβ burden extensively

Surprisingly, plaque burden in *ExN-Bace1*^*cKO*^*;AD* mice was reduced by 95-98% compared to controls (**Fig. 2g-k**), which was much more than anticipated given our findings in *OL-Bace1*^*cKO*^*;AD*. Plaque sizes were smaller and microgliosis was markedly reduced (**Extended Data Fig. 7**). Moreover, *ExN-Bace1*^*cKO*^*;AD* mice also showed a striking reduction in the amount of thalamic plaques. The unchanged levels of *Bace1* transcript in the thalamus of *ExN-Bace1*^*cKO*^*;AD* mice (**Extended Data Fig. 8a-b**) indicates that a significant amount of subcortical Aβ must be derived from cortico-thalamic axonal projections. Indeed, the inferior colliculus, receiving limited cortical input (**Extended Data Fig. 8c**) primarily from the auditory cortex^25^, was spared from plaque attenuation in *ExN-Bace1*^*cKO*^*;AD* mice.

### Deletion of *Bace1* in OLs and ExNs ablated Aβ burden in the entire cerebrum

To investigate if residual plaques found in *ExN-Bace1*^*cKO*^*;AD* hemibrains are primarily derived from OLs, we generated *Cnp-Cre Nex-Cre Bace1*^*fl/fl*^ *APP*^*NLGF*^ mice, hereby termed *OL-ExN-Bace1*^*cKO*^*;AD*. Indeed, *OL-ExN-Bace1*^*cKO*^*;AD* developed almost no plaques in the cerebrum (**Extended Data Fig. 9a-c**), indicating that OLs are major producers of Aβ *in vivo* besides ExNs. Nevertheless, the very few plaques present in the cortex of *OL-ExN-Bace1*^*cKO*^*;AD* mice hints at an ongoing, sub-optimal Aβ contribution from cells other than OLs and ExNs.

### Plaque deposition is non-linear to gene dosage of mutated humanized APP

It was nonetheless puzzling that plaque burden in the cortex and hippocampus was reduced by more than 95% in *ExN-Bace1*^*cKO*^*;AD* animals as we had expected that the residual plaque burden would reflect the contribution of OLs (30%). Both Aβ fibrillation *in vitro* and plaque formation *in vivo* follow sigmoidal growth kinetics^26,27^, and a threshold level of Aβ accumulation is a prerequisite for plaque seeding to occur. This threshold level apparently cannot be reached without the neuronal contribution. Fittingly, compared to homozygous *APP*^*NLGF*^ mice, heterozygotes did not develop 50% plaque burden, but rather less than 10% (**Extended Data Fig. 9d-e**). This highlights the non-linear relationship between APP processing or Aβ production and the resulting plaque load.

### Aβ concentration is non-linear to plaque deposition

Lastly, we performed a sensitive electroche-miluminescence assay for different Aβ species (Aβ38, 40, 42) to determine total Aβ levels. As inputs, we analyzed the soluble (Tris-NaCl) and insoluble (SDS-soluble; representing Aβ primarily bound in plaques^28^) fractions^29^ of microdissected cortex for gray matter and corpus callosum (CC) for WM (**Fig. 3a-b**). *OL-Bace1*^*cKO*^*;AD* brains contained less insoluble Aβ compared to controls, which was more apparent in the WM fraction. *OL-Bace1*^*cKO*^*;AD* brains also showed a concomitant decrease of soluble Aβ, which was stronger in WM. *ExN-Bace1*^*cKO*^*;AD* brains were almost devoid of insoluble Aβ much like our LSM plaque observation. However, there was still a moderate amount (16.459% ± 0.021) of soluble cortical Aβ in *ExN-Bace1*^*cKO*^*;AD* tissue extracts. In contrast to the significant soluble Aβ reduction in WM of *OL-Bace1*^*cKO*^*;AD*, the residual amount of Aβ was higher in the WM fraction from *ExN-Bace1*^*cKO*^*;AD* brains (32.282% ± 0.072). In short, we observed less soluble and insoluble Aβ in *OL-Bace1*^*cKO*^*;AD* brain extracts, mirroring the LSM data, and although plaque amount was marginally low in the *ExN-Bace1*^*cKO*^*;AD* brains, an adequate amount of soluble Aβ was still generated by non-ExN sources of Aβ, including OLs and potentially other cell types.

**Figure 3.**
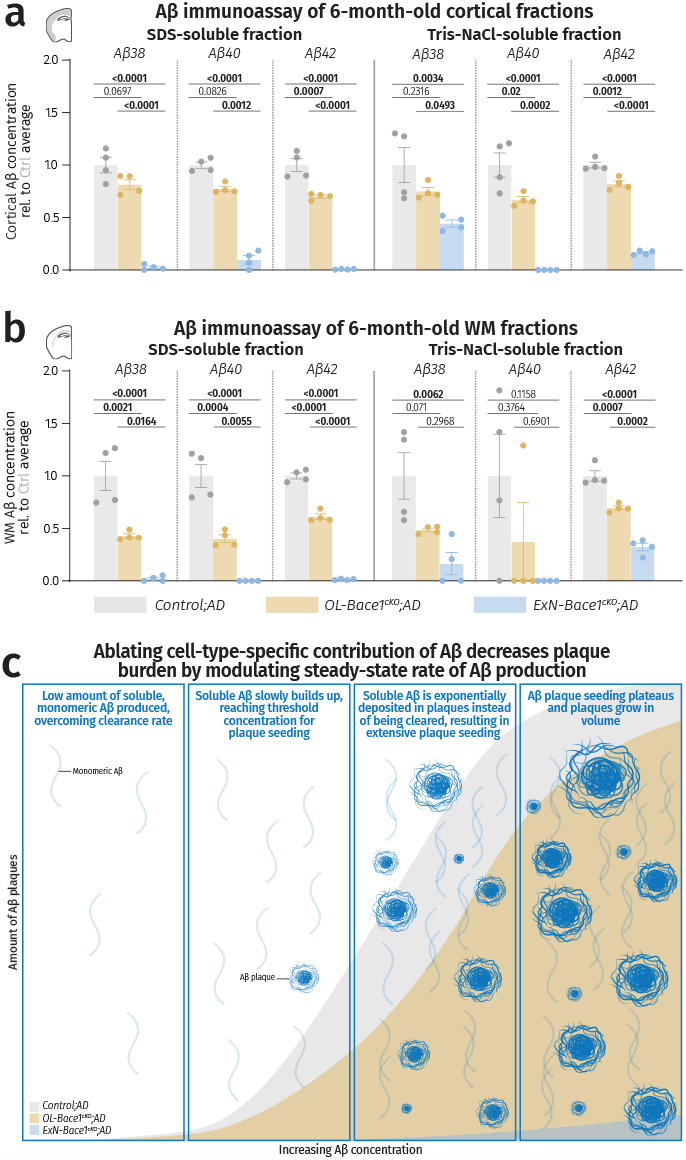
(a-b) Cell-type-specific deletion of *Bace1* alters levels of soluble and insoluble Aβ. Aβ electrochemiluminescence immunoassay data of insoluble (SDS-soluble, left) and soluble (Tris-NaCl-soluble, right) lysates of microdissected cortical (a) and WM tissues (b) from control and cKO fractions of 6-month-old male mouse hemibrains. Triplex immunoassay measured Aβ38, 40, and 42 and data points were normalized to *Control;AD* samples. Of note, SDS-soluble fractions from both regions mirrored LSM data, while Tris-NaCl-soluble fraction revealed a substantial amount of soluble Aβ still being produced, even in *ExN-Bace1*^*cKO*^*;AD* mice, signifying an residual Aβ production from other cells. Statistical analysis: one-way ANOVA with Tukey multiple comparison tests (p-values indicated in graphs with significance highlighted in bold). Bars represent means with SEM and individual data points displayed. Raw unnormalized data is available in **Extended Data Table 2. (c) Working model of modulating cell-type-specific Aβ contributions**. Selectively ablating Aβ from specific cell types results in steady-state rate change of Aβ production, causing exponentially slower plaque growth that follows a sigmoidal growth curve.

### Absence of myelin alterations upon *Bace1* cKO

We have recently shown that OL dysfunction drives neuronal amyloid deposition in AD mouse models^10^. To exclude this as a confounding factor in the present study, we compared myelin alteration in *OL-Bace1*^*cKO*^*;AD* and *ExN-Bace1*^*cKO*^*;AD* mice, but found no changes in myelin profiles or ultrastructure^30^ (**Extended Data Fig. 10**).

### OLs are active contributors to Aβ burden predominantly derived from ExNs

In conclusion, our work identifies OLs as contributors to Aβ load *in vivo* by using a second generation AD mouse model that expresses a humanized/mutated *APP* gene under its own regulatory elements. Our findings highlight that OLs, a glial cell type, are key players in AD – even in the context of establishing primary Aβ pathology. The OL-derived contribution to Aβ is most evident in regions where OLs predominate, such as in WM tracts. At the same time, we discovered that soluble Aβ is still moderately produced in *ExN-Bace1*^*cKO*^*;AD* brains by non-ExN sources, primarily OLs, but this amount of Aβ is insufficient to substantially seed plaques (**Fig. 3c**). More importantly, the detectable amounts of soluble Aβ in tissue extracts yet low plaque burden in *ExN-Bace1*^*cKO*^*;AD* brains together support a non-linear relationship between cellular Aβ production and plaque deposition. This was further exemplified by the presence of approximately 10% and not 50% of plaques in heterozygous *APP*^*NLGF*^ mice with 50% of mutated *APP* gene dosage. Taken together, a decrease in plaque burden does not linearly correlate to reduction in total Aβ production, highlighting the importance of stalling Aβ generation early enough to prevent reaching threshold Aβ levels for plaque deposition. It is likely that lower amounts of Aβ would lead to more efficient clearing in the brain, further reducing plaque seeding.

Finally, the high expression level of amyloidogenic pathway genes in OLs was contrasted by the smaller relative contribution to overall Aβ deposition, even within WM tracts. This could either be explained by the difference in number, localization, or size of neurons and OLs^31^. There is also the alternative possibility that Aβ processing is more efficient in neuronal compartments. We also demonstrate that cortical/ hippocampal projection neurons are the primary source of Aβ, not only locally, but also distally in subcortical projection areas. Beyond their abundance and neuronal activity^32^, however, what makes ExNs specifically so efficient at producing Aβ remains elusive. Further work is necessary to define the subcellular compartments of APP processing in OLs. Aβ isoforms could also differ between OLs and ExNs. Identifying the mechanisms that slow down Aβ generation in OLs, despite the abundance of processing substrate and enzymes, could pave the way for novel therapies targeting Aβ generation. Our findings are also relevant for therapeutic strategies with BACE1 inhibitors, which as suggested^33,34^ may have a potential in preventing amyloidosis before threshold levels are reached.

## Extended data figures

**Extended Data Figure 1.**
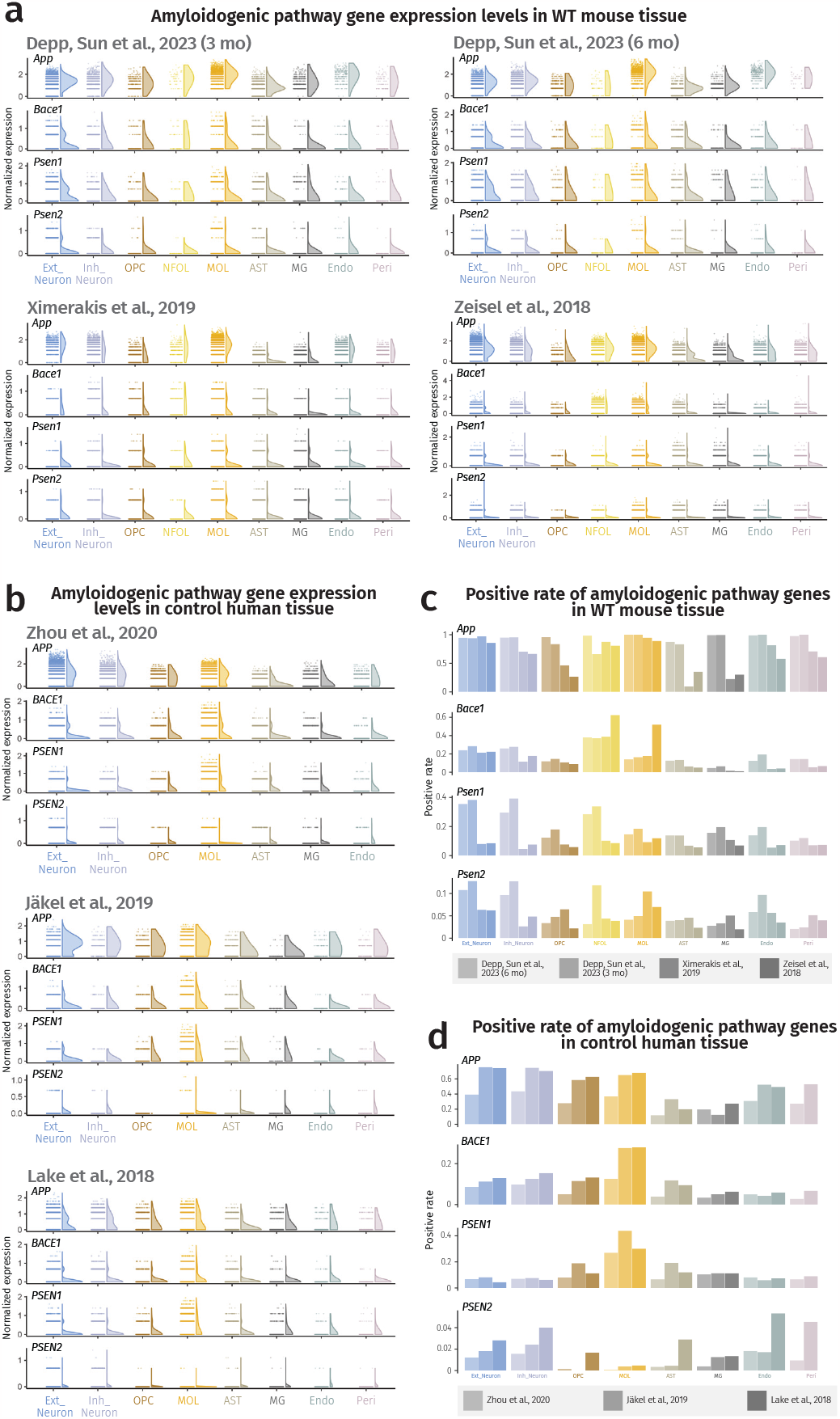
Expression levels and positive rates of amyloidogenic pathway genes across all cell types in the nervous system. (**a**) Expression level half violin plots of amyloidogenic pathway genes in all cell types of mouse nervous tissue inputs with SCTransform normalization method from all chosen mouse datasets^10-12^. (**b**) Expression level half violin plots of amyloidogenic pathway genes in all cell types of human nervous tissue inputs with SCTransform normalization from all chosen human datasets^13-15^. (**a, b**) Half violins represent aggregated expression levels of respective genes from each cell type and data points refer to individual expression levels from single cells or nuclei normalized by the SCTransform method. (**c**) Positive rate barplots of amyloidogenic pathway genes in all cell types of mouse nervous tissue inputs from all chosen mouse datasets^10-12^. (**d**) Positive rate barplots of amyloidogenic pathway genes in all cell types of human nervous tissue inputs with SCTransform normalization method from all analyzed human datasets^13-15^. The results published here are based on data obtained from GEO and the AD Knowledge Portal.

**Extended Data Figure 2.**
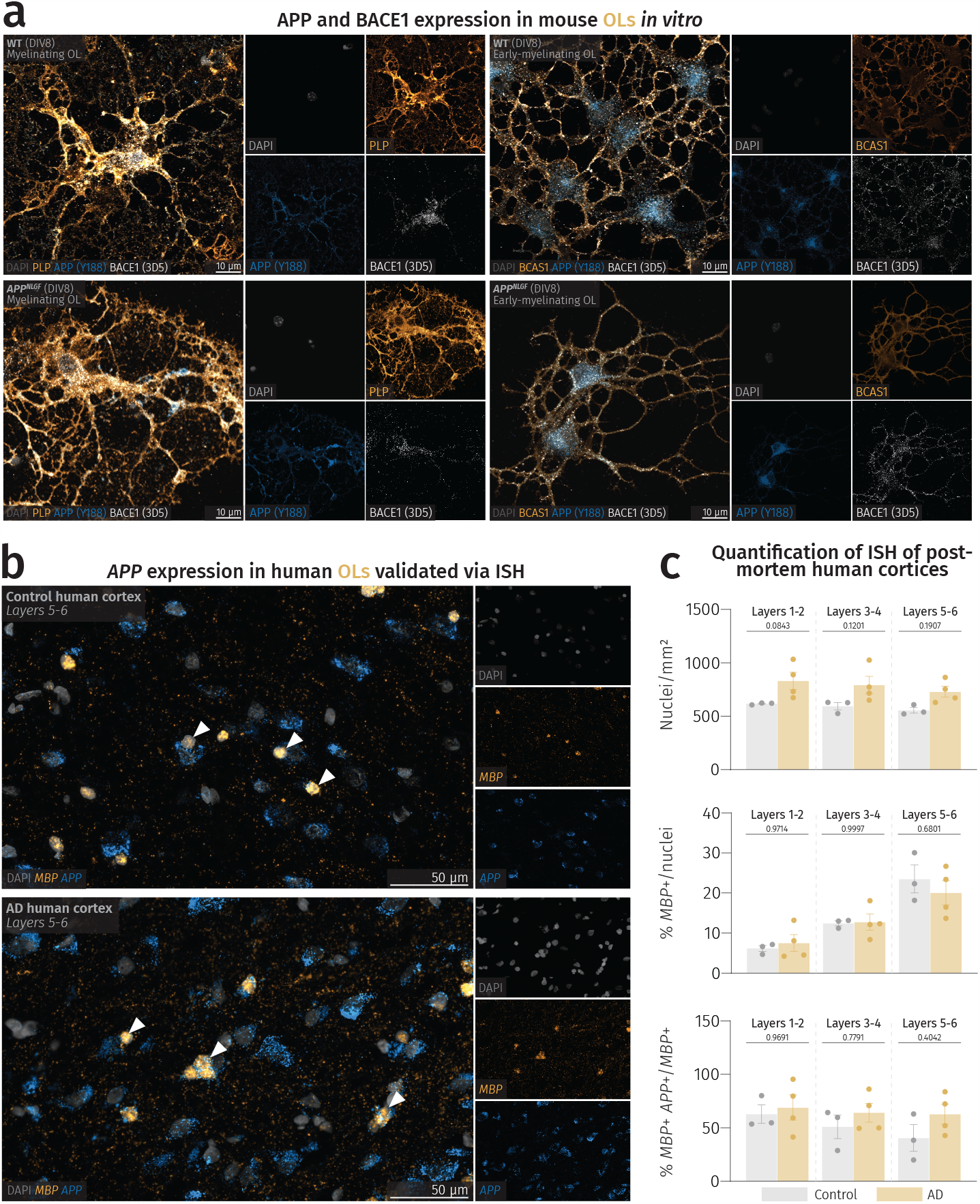
Expression of amyloidogenic pathway proteins and transcripts in mouse and human OLs. (**a**) Confocal images of OL *in vitro* cultures with APP and BACE1 present in the soma and cellular processes of both myelinating (left) and early-myelinating (right) OLs. Localization of both proteins appear unchanged between OLs cultured from WT (top) and *APP*^*NLGF*^ (bottom) mice. (**b**) ISH confocal images of human cortical layers 5-6 with visible *APP* puncta in *MBP*+ cells of control (top) and AD (bottom) patients. Arrowheads point to *APP*-expressing OLs. (**c**) Quantification of nuclear count, *MBP*+ nuclei, and *MBP*+ *APP*+ nuclei in control (n=3) and AD (n=4) patients. One-way ANOVA was performed with Sidak multiple comparison tests (p-values indicated in graphs with significance highlighted in bold) comparing AD patients to controls. Bars represent means with SEM with all data points displayed.

**Extended Data Figure 3.**
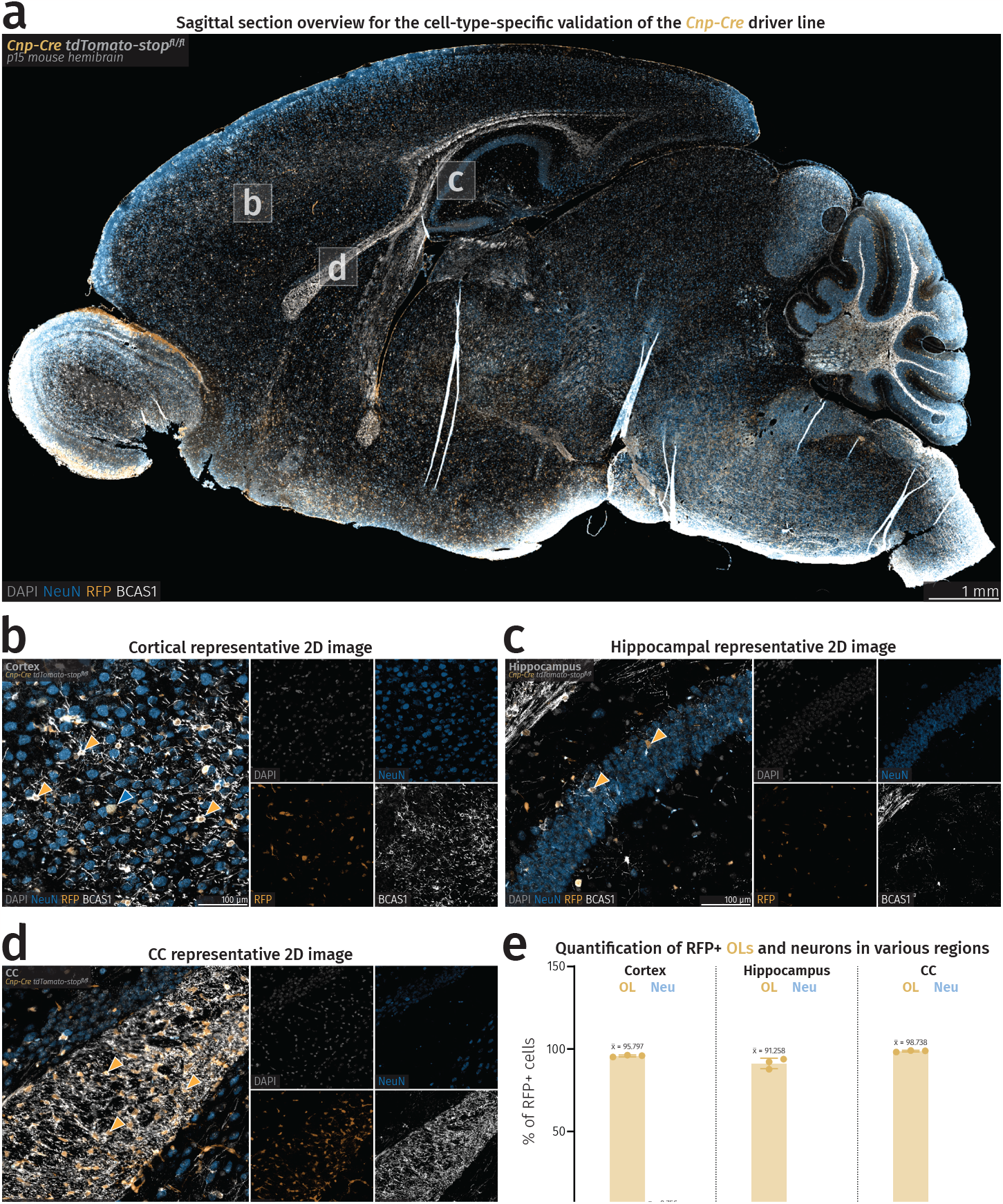
Validation of *Cre* specificity in *Cnp-Cre* stop flox tdTomato mice. (**a**) Fluorescence microscopy sagittal overview of a *Cnp-Cre* stop flox tdTomato mouse. (**b-d**) Closeup images of cortex, hippocampus, and CC of a *Cnp-Cre* stop flox tdTomato mouse. Yellow arrowheads point to RFP+ OLs and the blue arrowhead points to a single RFP+ neuron in the cortex. (**e**) Barplots showing percentages of RFP+ OLs and neurons in specific brain regions. Mean percentage values are shown above each bar. Rounded average total number of cells considered for quantification is as follows: Cortex-OLs=283, cortex-neurons=9232, hippocampus-OLs=66, hippocampus-neurons=3203, CC-OLs=344, CC-neurons=1.

**Extended Data Figure 4.**
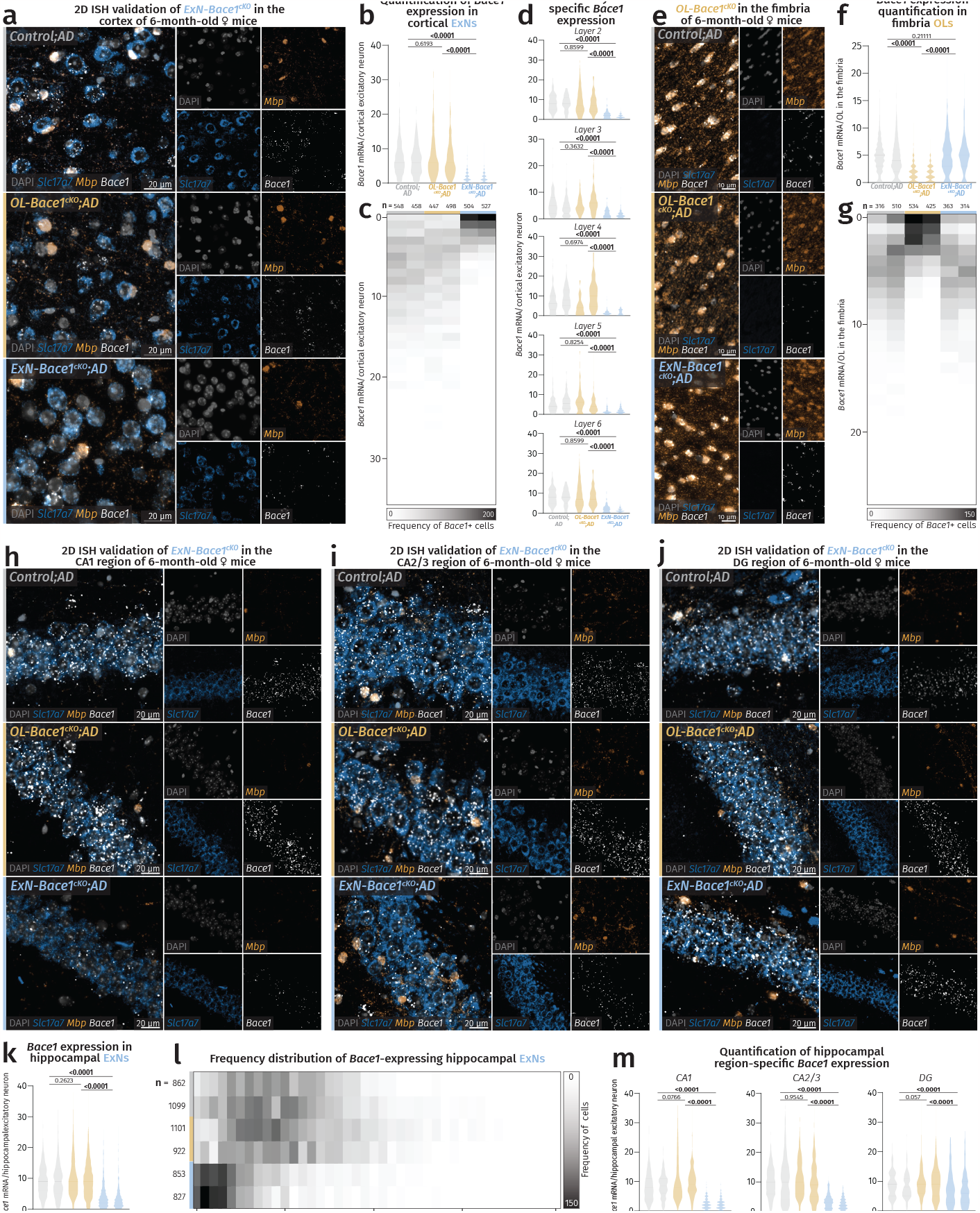
ISH validation of *Bace1* cKO in OL-*Bace1* ^*KO*^*;AD* and ExN-*Bace1*^*cKO*^*;AD*. (**a-d**) ISH validation of *Bace1* cKO in cortical ExNs. (**e-g**) ISH validation of *Bace1* cKO in fimbria OLs. (**h-m**) ISH validation of *Bace1* cKO in hippocampal ExNs. (**a**) Fluorescence microscopy images of cortices showing reductions of *Bace1* transcripts in ExNs of *ExN-Bace1*^*cKO*^*;AD* samples. (**b**) Violin plots showing individual mouse distribution of *Bace1* puncta/cortical ExN nuclei. (**c**) Frequency distribution heatmap of individual mouse distribution of *Bace1* puncta/cortical ExN nuclei. (**d**) Violin plots showing cortical layer-specific individual mouse distribution of *Bace1* puncta/ExN nuclei. (**e**) Fluorescence microscopy images of hippocampal fimbriae showing reductions of *Bace1* transcripts in OLs of *OL-Bace1*^*cKO*^*;AD* samples. (**f**) Violin plots showing individual mouse distribution of *Bace1* puncta/fimbria OL nuclei. (**g**) Frequency distribution heatmap of individual mouse distribution of *Bace1* puncta/fimbria OL nuclei. (**h-j**) Fluorescence microscopy images of hippocampal CA1, CA2/3, and DG respectively showing reductions of *Bace1* transcripts only in ExNs of *ExN-Bace1*^*cKO*^*;AD* samples. (**k**) Violin plots showing individual mouse distribution of *Bace1* puncta/hippocampal ExN nuclei. (**l**) Frequency distribution heatmap of individual mouse distribution of *Bace1* puncta/hippocampal ExN nuclei. (**m**) Violin plots showing hippocampal region-specific individual mouse distribution of *Bace1* puncta/ExN nuclei. For (**b**,**d**,**f**,**k**,**m**), Kruskal-Wallis test was performed alongside multiple comparison tests with Dunn’s correction (p-values indicated in graphs with significance highlighted in bold) comparing aggregated data of control, *OL-Bace1*^*cKO*^*;AD*, and *ExN-Bace1*^*cKO*^*;AD* mice (n=2/group). Solid lines represent median and faded lines represent quartiles. For (**c**,**g**,**l**), n-numbers refer to amount of nuclei considered for each region and cell type analysis and make up the data cloud for violin plots shown in (**b**,**d**,**f**,**k**,**m**).

**Extended Data Figure 5.**
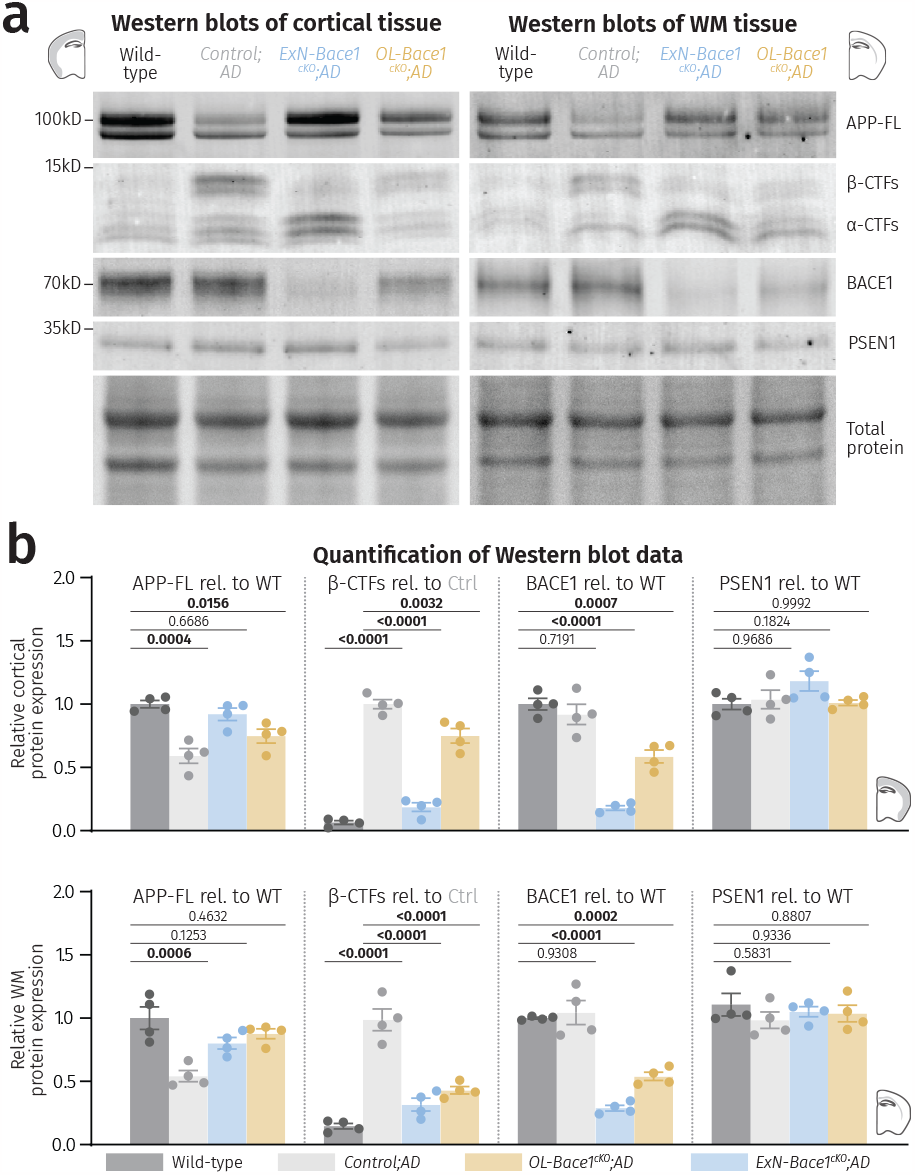
Cell-type-specific deletion of *Bace1* alters APP processing. (**a**) Fluorescent immunoblots and total protein content of microdissected cortical and WM tissues targeting key amyloidogenic proteins in insoluble lysates. (**b**) Immunoblot quantification showing APP processing in WT, control, *ExN-Bace1*^*cKO*^*;AD*, and *OL-Bace1*^*cKO*^*;AD* (n=4/group) insoluble lysates. Top-cortical, bottom-WM. All immunoblots were normalized to WT relative protein amount except β-CTFs which were normalized to control *AD* relative protein amount. Data was statistically analyzed via one-way ANOVA was performed with Tukey multiple comparison tests (p-values indicated in graphs with significance highlighted in bold). Bars represent means with SEM and individual data points displayed.

**Extended Data Figure 6.**
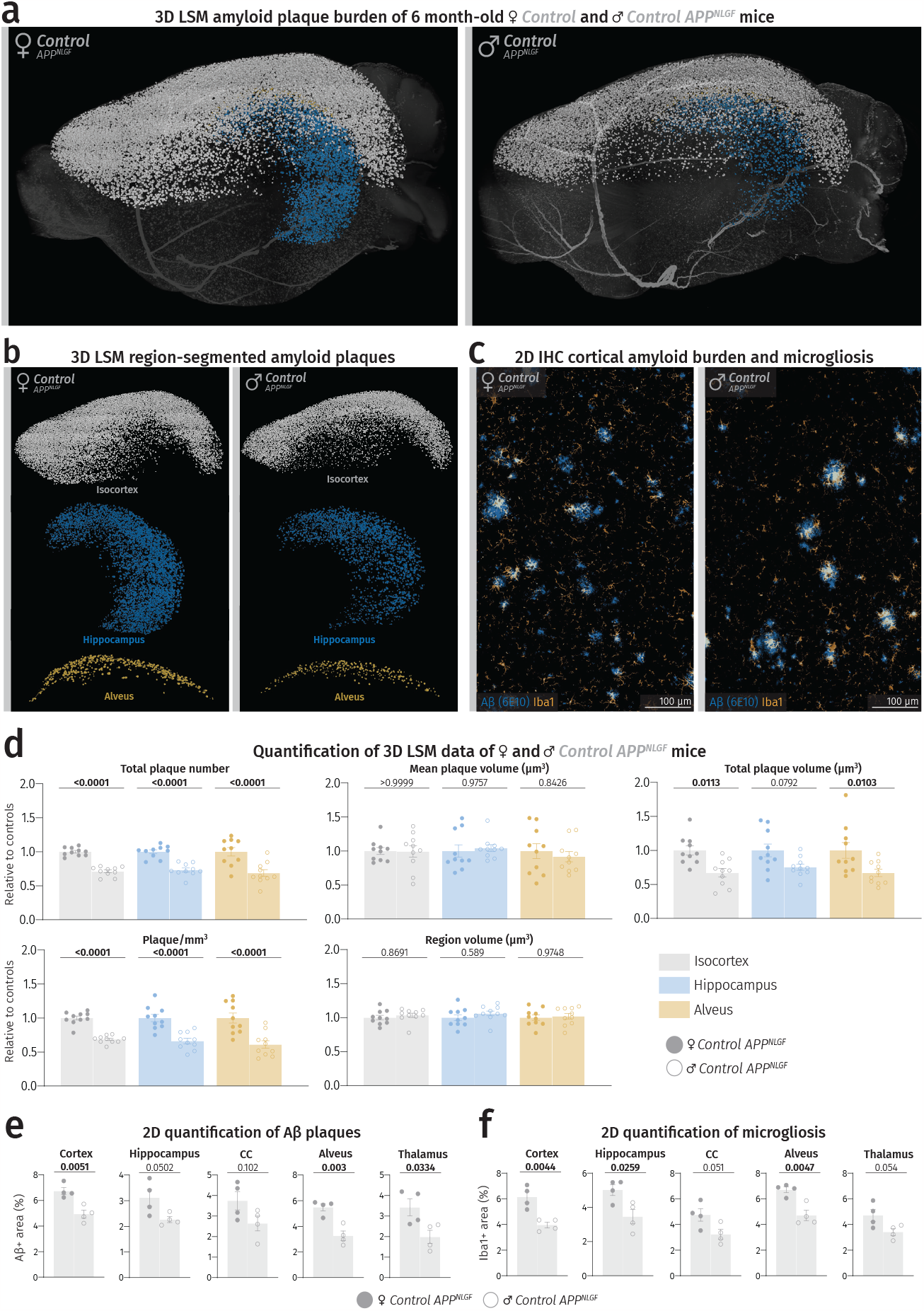
Female *APP*^*NLGF*^ animals develop more Aβ plaque burden compared to age-matched male *APP*^*NLGF*^ animals. (**a**) LSM 3D visualization of female and male control *APP*^*NLGF*^ hemibrains at 6 months of age. (**b**) Brain region-segmented plaques of female and male control *APP*^*NLGF*^ hemibrains. Color-region allocation is as follows: White-isocortex, blue-hippocampus, yellow-alveus. (**c**) Fluorescence microscopy images of female and male control *APP*^*NLGF*^ cortices. (**d**) Quantification of LSM data between female and male control *APP*^*NLGF*^ hemibrains (n=10/sex). Male data points were normalized to female data. Filled shapes represent male and hollowed shapes represent female mice. For each parameter, unpaired, two-tailed Student’s t-test was performed (p-values indicated in graphs) comparing males to females. Bars represent means with SEM and individual data points displayed. (**e**,**f**) Quantification of Aβ load and microgliosis between male and female control *APP*^*NLGF*^ mice spanning different regions. Unpaired, two-tailed Student’s t-test was performed for each regional quantification (p-values indicated in graphs with significance highlighted in bold) comparing males to females. Bars represent means with SEM and individual data points displayed. Raw unnormalized data is available in **Extended Data Table 1**.

**Extended Data Figure 7.**
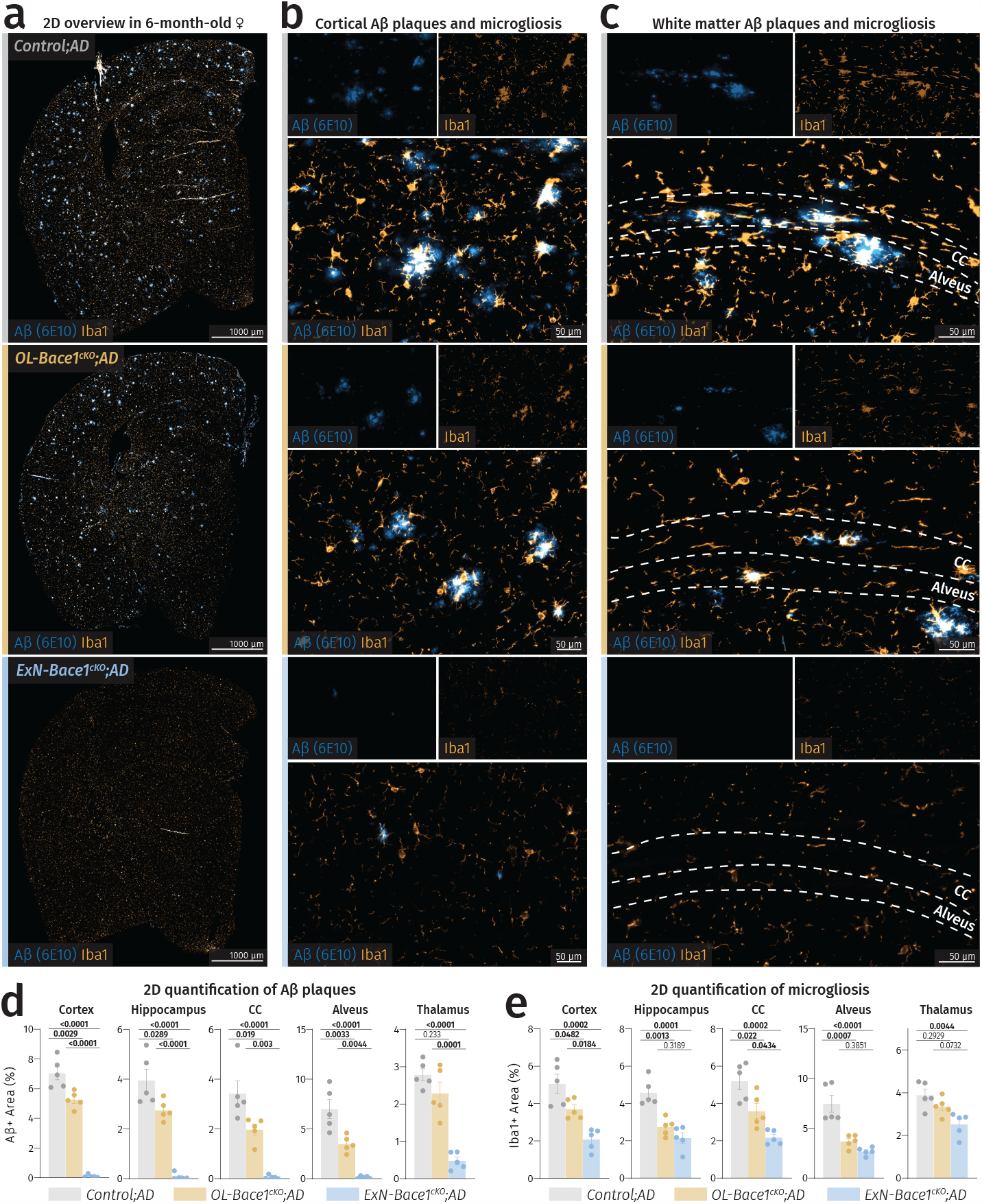
Microgliosis is proportional to Aβ burden. (**a**) Coronal sections of female control, *OL-Bace1*^*cKO*^*;AD*, and *ExN-Bace1*^*cKO*^*;AD* mouse hemibrains stained for microglia (Iba1) and Aβ (6E10). (**b**,**c**) Closeup images of cortex and WM of control and cKO mice showing moderate and marked reductions of both Aβ deposits and microgliosis in *OL-Bace1*^*cKO*^*;AD* and *ExN-Bace1*^*cKO*^*;AD* samples respectively. Inherent changes in microgliosis could thus be excluded as microglia only appear reactive to plaques and not in regions devoid of them. (**d**,**e**) Quantification of Aβ load and microgliosis between controls and cKOs spanning different regions. Microgliosis was shown to be directly linked to plaque load. One-way ANOVA was performed with Tukey multiple comparison tests (p-values indicated in graphs with significance highlighted in bold). Bars represent means with SEM and individual data points displayed.

**Extended Data Figure 8.**
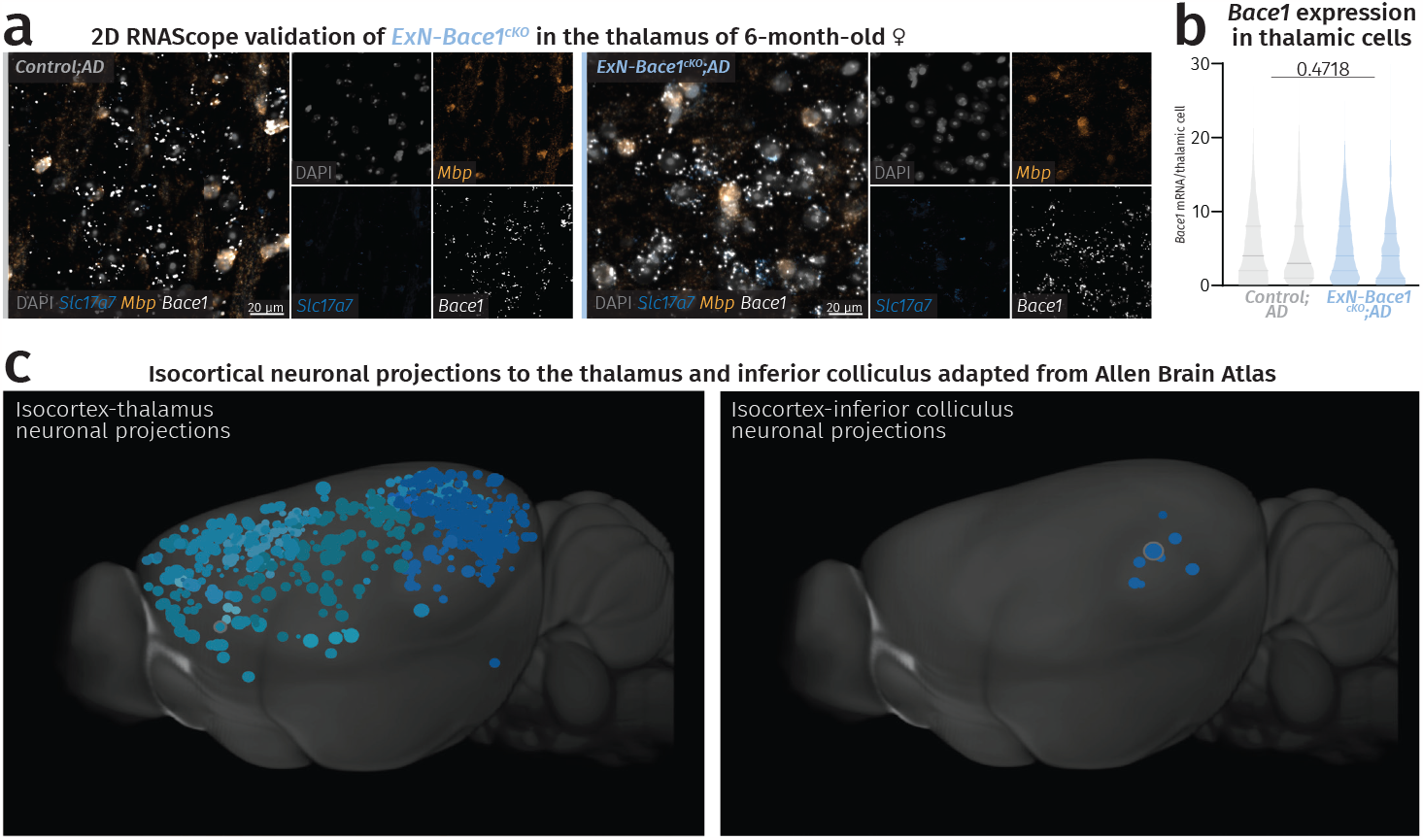
(a-b) The thalamus is not a *Nex-Cre* recombination territory in *ExN-Bace1*^*cKO*^*;AD mice*. (**a**) Fluorescence microscopy images of thalami of control and *ExN-Bace1*^*cKO*^*;AD* samples with no apparent reduction in *Bace1* transcripts in ExNs. (**b**) Violin plots showing individual mouse distribution of *Bace1* puncta/thalamic nuclei. Mann-Whitney test was performed (p-values indicated in graphs) comparing aggregated data of control and *ExN-Bace1*^*cKO*^*;AD* mice (n=2/group), whereby no significant difference was measured. Solid lines represent median and faded lines represent quartiles. **(c) The thalamus but not the inferior colliculus receives ample cortical input**. Isocortical regions containing neuronal projections into the thalamus (left) and the inferior colliculus (right). Inferior colliculus primarily receives cortical input from the auditory cortex. Images were adapted from the Allen Brain Atlas: Mouse Connectivity: Projection (connectivity.brain-map.org/).

**Extended Data Figure 9.**
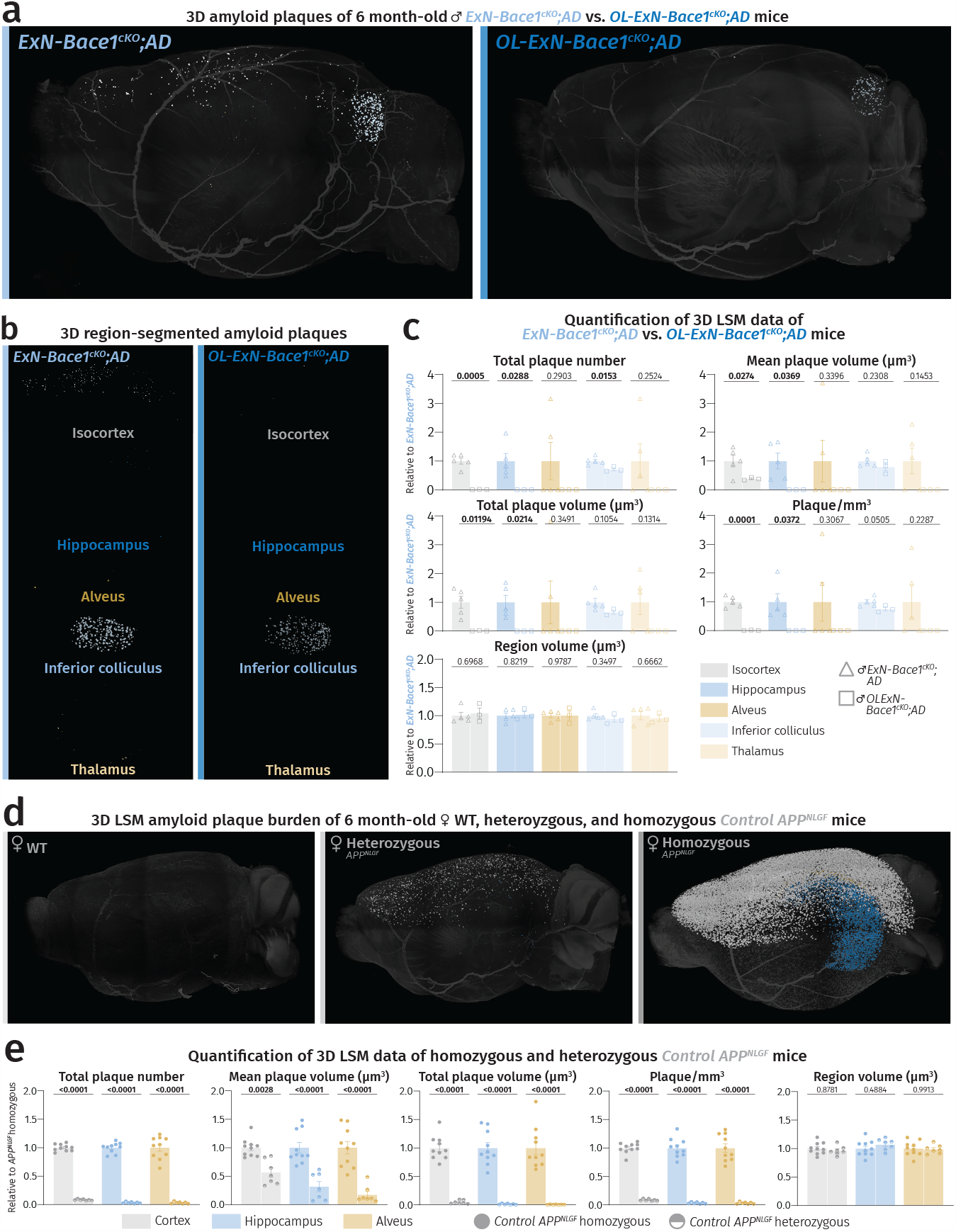
(a-c) Double *Bace1* cKO in OLs and ExNs ablated cerebral Aβ burden. Light sheet microscopy data of plaque burden (Congo red) comparing 6-month-old *OL-ExN-Bace1*^*cKO*^*;AD* male mice to age- and sex-matched *ExN-Bace1*^*cKO*^*;AD* mice. Color-region allocation is as follows: White-isocortex, blue-hippocampus, yellow-alveus, pastel blue-inferior colliculus, pastel yellow-thalamus. (**a**) LSM 3D visualization of *ExN-Bace1*^*cKO*^*;AD* and *OL-ExN-Bace1*^*cKO*^*;AD* male mouse hemibrains. (**b**) Brain region-segmented plaques of *ExN-Bace1*^*cKO*^*;AD* and *OL-ExN-Bace1*^*cKO*^*;AD* male mouse hemibrains. (**c**) Quantification of LSM data between *ExN-Bace1*^*cKO*^*;AD* (n=5) and *OL-ExN-Bace1*^*cKO*^*;AD* (n=3) hemibrains. Normalization of *OL-ExN-Bace1*^*cKO*^*;AD* data points was performed to *ExN-Bace1*^*cKO*^*;AD*. Hollowed triangles represent *ExN-Bace1*^*cKO*^*;AD* and hollowed squares represent *OL-ExN-Bace1*^*cKO*^*;AD*. **(d-e) Plaque deposition is not directly proportional to *APP***^***NLGF***^ **gene dosage**. (**d**) LSM 3D visualization of female WT, heterozygous, and homozygous *APP*^*NLGF*^ hemibrains at 6 months of age. (**e**) Quantification of LSM data between female homozygous (n=10) and heterozygous *APP*^*NLGF*^ mice (n=7). Heterozygous data points were normalized to homozygous data. Circles represent homozygous *APP*^*NLGF*^ mice and half-filled circles represent heterozygous *APP*^*NLGF*^ mice. (**c**,**e**) For each parameter, unpaired, two-tailed Student’s t-test was performed (p-values indicated in graphs) comparing the two groups. Bars represent means with SEM and individual data points displayed. Raw unnormalized data is available in **Extended Data Table 1**.

**Extended Data Figure 10.**
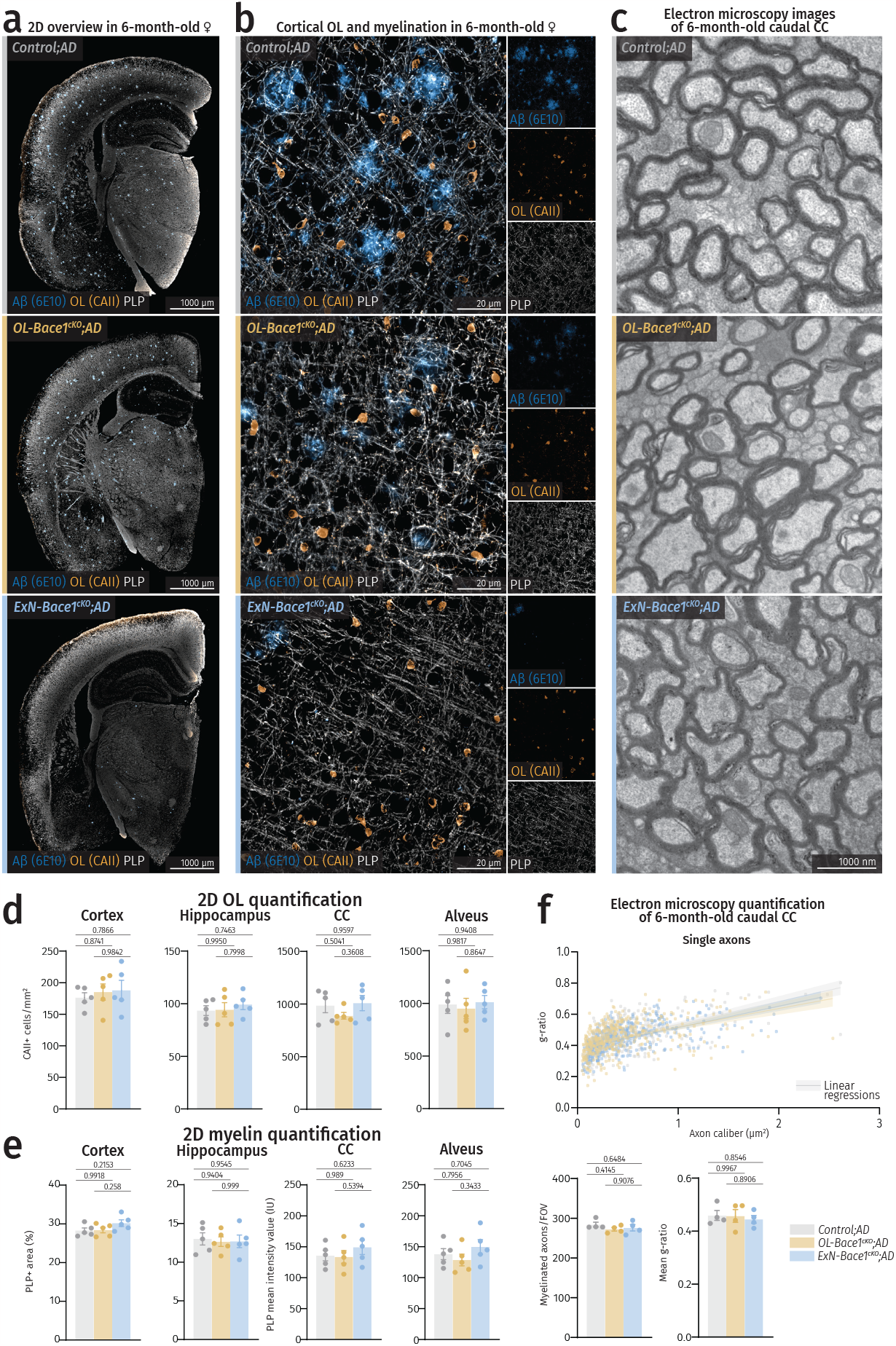
*Bace1* cKO does not alter myelination profile. (**a**) Coronal section overviews of control, *OL-Bace1*^*cKO*^*;AD*, and *ExN-Bace1*^*cKO*^*;AD* hemibrains stained for Ols (CAII) and myelin (PLP). (**b**) Closeups of fluorescence microscopy images of cortices of controls and cKOs highlighting unchanged density of OLs and myelination in both cKO lines at a gross level. (**c**) Representative electron micrographs of caudal corpus callosum (CC) of controls and cKOs at 6 months of age. (**d**) Quantification of OL density between controls and cKOs (n=5/group) spanning different regions. (**e**) Quantification of myelin density between controls and cKOs (n=5/group) spanning different regions. As CC and alveus are densely myelinated tracts, mean intensity values were instead measured. (**f**) Analysis of myelin thickness via g-ratio measurement with single dots representing single myelinated axons quantified (*Control;AD*=397, *OL-Bace1*^*cKO*^*;AD*=417, *ExN-Bace1*^*cKO*^*;AD*=394). Lines represent linear regressions of each group and shaded area indicates error bars. (**g**) Myelinated axon counts and mean g-ratio comparisons from electron micrographs of controls and cKOs (n=4/group). For (**d**,**e**,**g**), one-way ANOVA was performed with Tukey multiple comparison tests (p-values indicated in graphs with significance highlighted in bold). Bars represent means with SEM and individual data points displayed.

**Extended Data Table 1.**
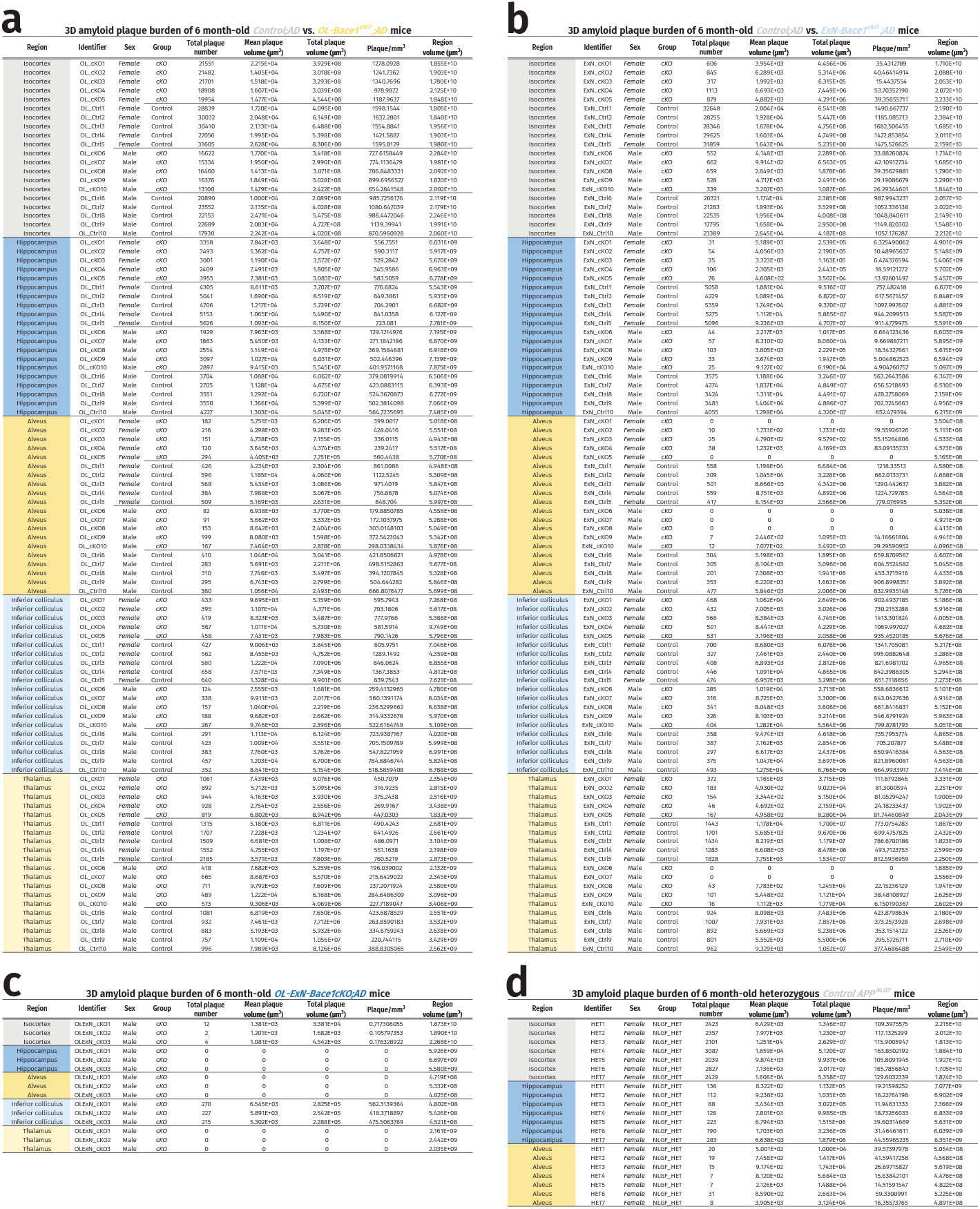
Raw LSM data of all hemibrains pertaining to this study. (**a**) Raw LSM data of control (n=5/sex) and *OL-Bace1*^*cKO*^*;AD* (n=5/ sex) hemibrains at 6 months of age (**Figure 2f**). (**b**) Raw LSM data of control (n=5/sex) and *ExN-Bace1*^*cKO*^*;AD* (n=5/sex) hemibrains at 6 months of age (**Figure 2k**). (**d**) Raw LSM data of female heterozygous control *APP*^*NLGF*^ (n=7) hemibrains at 6 months of age (**Extended Data Figure 9e**).

**Extended Data Table 2.** Raw Aβ immunoassay data pertaining to this study. Normalized Aβ immunoassay data of control, *OL-Bace1*^*cKO*^*;AD*, and *ExN-Bace1*^*cKO*^*;AD* (n=4/group) at 6 months of age (**Fig. 3**). Cortical and WM data are both represented with adjacent columns containing Aβ38, Aβ40, and Aβ42 values. Top half represents measurements of the SDS-soluble fractions and bottom half represents measurements of the Tris-NaCl-soluble fractions. All values below the detection range of the immunoassay were reported as 0.

| Identifier | Fraction | Cortex |  |  | White Matter |  |  |
| --- | --- | --- | --- | --- | --- | --- | --- |
| | | Aβ38 (pg/mg) | Aβ40 (pg/mg) | Aβ42 (pg/mg) | Aβ38 (pg/mg) | Aβ40 (pg/mg) | Aβ42 (pg/mg) |
| Control1 | SDS | 3098.850827 | 296.6003816 | 22041.12568 | 2263.555768 | 176.7611396 | 12828.69251 |
| Control2 | SDS | 3597.00014 | 300.0696096 | 23360.44319 | 2338.565402 | 170.9270572 | 12453.96428 |
| Control3 | SDS | 4381.077278 | 336.6209498 | 18531.45301 | 3836.431054 | 260.4572027 | 14050.61227 |
| Control4 | SDS | 4049.397745 | 324.5622748 | 18340.93248 | 3697.391148 | 251.3667413 | 13716.79943 |
| OL_cKO1 | SDS | 3383.003919 | 265.548933 | 14917.34181 | 1202.176776 | 73.44641268 | 7668.487559 |
| OL_cKO2 | SDS | 3378.635441 | 239.1921467 | 13563.0966 | 1247.436655 | 74.29861336 | 7708.640423 |
| OL_cKO3 | SDS | 2711.040864 | 234.2251699 | 14464.4998 | 1484.3957 | 105.3259203 | 9140.198276 |
| OL_cKO4 | SDS | 2855.146369 | 235.6212532 | 14717.88493 | 1256.247821 | 93.50412783 | 7948.067807 |
| ExN_cKO1 | SDS | 218.6080663 | 43.16058151 | 79.47845463 | 0 | 0 | 93.57319751 |
| ExN_cKO2 | SDS | 49.81558523 | 21.91478309 | 206.2281853 | 86.91726698 | 0 | 223.2603588 |
| ExN_cKO3 | SDS | 0 | 58.09359015 | 89.7747642 | 0 | 0 | 103.4913095 |
| ExN_cKO4 | SDS | 100.251635 | 0 | 256.4144541 | 160.2461865 | 0 | 293.4330212 |
| Control1 | Tris-NaCl | 206.2626946 | 18.24124215 | 165.6060914 | 106.4284342 | 2.163312874 | 139.5841322 |
| Control2 | Tris-NaCl | 190.1284236 | 15.08643851 | 147.9161636 | 93.40120174 | 4.003693224 | 113.8610996 |
| Control3 | Tris-NaCl | 362.0259044 | 24.31185089 | 150.5918218 | 217.4422531 | 5.120354785 | 115.9770306 |
| Control4 | Tris-NaCl | 354.6724012 | 24.92636741 | 148.5821138 | 228.9308581 | 0 | 116.7196294 |
| OL_cKO1 | Tris-NaCl | 192.0970948 | 13.6869043 | 120.933969 | 83.20744112 | 4.208642658 | 83.65571051 |
| OL_cKO2 | Tris-NaCl | 203.4183371 | 12.62893911 | 127.3558341 | 82.77732594 | 0 | 84.08408588 |
| OL_cKO3 | Tris-NaCl | 199.4401724 | 13.52153311 | 117.248305 | 72.11099676 | 0 | 90.62884652 |
| OL_cKO4 | Tris-NaCl | 238.8329797 | 15.51804572 | 137.4136787 | 74.71811105 | 0 | 79.18581242 |
| ExN_cKO1 | Tris-NaCl | 113.1209959 | 0 | 21.98538648 | 72.14463275 | 0 | 47.02577122 |
| ExN_cKO2 | Tris-NaCl | 144.0615627 | 0 | 28.44308708 | 0 | 0 | 26.98032503 |
| ExN_cKO3 | Tris-NaCl | 103.1649565 | 0 | 23.03873201 | 0 | 0 | 39.52224016 |
| ExN_cKO4 | Tris-NaCl | 133.5910327 | 0 | 27.38189382 | 33.60601943 | 0 | 43.40826879 |

## Materials and methods

### Reanalysis of snRNA-Seq and scRNA-Seq data from mouse and human nervous system

External single-cell/nuclei transcriptome sequencing (snRNA-Seq/scRNA-Seq) datasets were collected and screened for expressions of *APP, BACE1, PSEN1, PSEN2, ADAM10, ADAMTS4*, and *MEP1B* across major cell populations in the CNS. In total, four mouse datasets from Depp et al., 2023 (GSE178295, GSE208683)^10^, Ximerakis et al., 2019 (GSE129788)^11^, and Zeisel et al., 2018 (SRP135960)^12^, and three human datasets from Zhou et al., 2020 (access via AD Knowledge Portal under study snRNAseqAD_TREM2)^13^, Jäkel et al., 2019 (GSE118257) ^14^, and Lake et al., 2018 (GSE97942)^15^. All data was processed with R package Seurat (V.4.3.0)^35^ based on protocols published in the original study. Cell type annotations are crosschecked with cluster-specific gene signatures. Afterwards, major CNS cell populations including excitatory neuron (Ext_Neuron), inhibitory neuron (Inh_Neuron), oligodendrocyte precursor cell (OPC), newly formed oligodendrocyte (NFOL), mature oligodendrocyte (MOL), astrocyte (AST), microglia (MG), endothelial cells (Endo), and pericyte are subset for further screening APP metabolism related gene expressions. Each subset dataset underwent renormalisation, high variable feature calculation, and scaling using SCTransform pipeline with default parameters. Gene expression levels are visualized in half violin plots using R package raincloudplots^36^. Positive expression rate of each gene is calculated upon more than 1 UMI, and the relative proportion is visualized using R package ggplot2^37^.

### Mouse models, husbandry, and genotyping

All animal experiments were conducted in concordance with German animal welfare practices and local authorities. Mice were group-housed in the animal facility of Max Planck Institute for Multidisciplinary Sciences (MPI-NAT), City Campus with *ad libitum* food and regular cage maintenance. All mice were kept under a 12 h dark and 12 h light cycle. All animals are characterized as unburdened and only organ collection experiments were performed. Mouse strains were kept on a C57BL/6 background and both sexes were used throughout the study as sex dimorphism was evident in one of the primary models. The following mouse strains were utilized: *APP*^*NLGF16*^, *Bace1*^*fl/fl17*^, *Cnp-Cre*^*18*^, *Nex-Cre*^*19*^, and stop-flox tdTomato^20^. The crossbreeds generated and analyzed are the following: *Cnp-Cre Bace1*^*fl/fl*^ *APP*^*NLGF*^ to assess OL-Aβ contribution (*OL-Bace1*^*cKO*^*;AD*), *Nex-Cre Bace1*^*fl/fl*^ *APP*^*NLGF*^ (*ExN-Bace1*^*cKO*^*;AD*) to assess ExN-Aβ contribution, and *Cnp-Cre* stop flox tdTomato to validate *Cnp-Cre* specificity. Ages of animals analyzed are listed on the respective figures. Genotyping was carried out on ear clips from the marking process of individual animals (see individual strain references for genotyping protocols). For validation of genotypes, re-genotyping was performed on a small tail biopsy gathered after mice were euthanized prior to sample collection.

### Mouse tissue extraction

To acquire samples for imaging experiments, animals were euthanized and immediately flushed with cold phosphate buffered saline (PBS) until the liver was decolorized. Extracted tissues underwent immersive fixation in 4% paraformaldehyde (PFA) in 0.1 M phosphate buffer as no detrimental effects from immersive fixation compared to perfusion fixation was evident and previously reported^38^. To ensure sample integrity for electron microscopy, perfusion was done with 4% PFA, 2.5% glutaraldehyde in 0.1 M phosphate buffer (K+S buffer) after flushing with PBS. After 24 h, fixed tissues were washed and subsequently stored at 4°C long-term in PBS with the exception of samples for electron microscopy analysis which were stored at 4°C long-term in 1% PFA in PBS. For biochemical experiments, animals were sacrificed by cervical dislocation and their brains were quickly extracted before a quick wash in PBS. Tissues were then placed into a custom 1 mm-spaced coronal brain matrix developed in-house (Workshop, MPI-NAT, City Campus) for manual, sequential sectioning with blades. Brain slices were transferred to a glass plate on ice prior to microdissection of cortices, corpus callosa (CC), and hippocampi. Dissected tissues were snap-frozen and placed in -80°C until further use.

### Sample preparation and staining for LSM

Fixed tissues were processed for light-sheet microscopy (LSM) imaging using a modified iDISCO protocol as we previously reported^10^. Hemibrains were first transferred into 2 ml tubes and subjected to an ascending methanol wash in PBS (50%, 80%, 100% twice, 1 h each). To quench autofluorescence, we placed hemibrains overnight at 4°C in a 1:1:4 mixture of dimethyl sulfoxide (DMSO), 30% H_2_O_2_, and 100% methanol. We further dehydrated the samples by placing them in 100% methanol for 30 min at 4°C, 3 h at -20°C, and overnight storage at 4°C. The next day, we replaced the solution with 20% DMSO in methanol for 2 h before performing a descending methanol wash in PBS (80%, 50%, 0%, 1 h each) and a pre-permeabilization step with 0.2% Triton X-100 in PBS for 2 h. The samples were then permeabilized overnight at 37°C in a mixture containing 20% DMSO, 0.2% Triton X-100, and 22.5 mg/ml glycine dissolved in PBS. Samples were washed with a PBS solution with 0.2% Tween-20, 10 mg/ml heparin, and 5 mM sodium azide (PTwH, 2 h) prior to a 72 h incubation at 37°C in a 0.01% Congo red dye solution in PTwH. Congo red stock is kept at 0.5% w/v in 50% ethanol. Following labeling, hemibrains were washed in PTwH thrice for 10 min each prior to being subjected to a final ascending methanol wash in PBS (20%, 40%, 60%, 80%, 100%, 1 h each and overnight incubation in a 1:2 mixture of 100% methanol and dichloromethane (DCM). Finally, the samples were placed in 100% DCM for 1 h 40 min before clearing in ethyl cinnamate (ECI) for imaging. LSM imaging was performed in a sample chamber holding ECI. Of note, some cleared samples which had undergone LSM imaging could be made opaque by immersion in 100% methanol for 1 h and subjected to paraffin embedding before tissue sectioning to generate 2D slices. All incubation steps were done at RT unless stated otherwise.

### *In toto* LSM imaging and analysis

Cleared hemibrains were imaged with a LSM setup (UltraMicroscope II, LaVision Biotec) with a corrected dipping cap at 2x objective lens magnification. To acquire the images from the sagittal orientation, hemibrains were mounted with the medial side facing down in the sample chamber while submerged in ECI. The ImspectorPro (v.7.124, LaVision Biotec) software was used to visualize the samples in the mosaic acquisition mode with the following settings: 5 μm light sheet thickness, 20% sheet width, 0.154 sheet numerical aperture, 4 μm z-step size, 2,150 × 2,150 pixels field of view, dynamic focus steps of 5, dual light sheet illumination, and 100 ms camera exposure time. Red fluorescence of Congo red-stained hemibrains was recorded with 561 nm laser excitation at 80% laser power and a 585/40 nm emission filter.

Image stacks were imported and stitched with Vision4D (v3.2, Arivis). Regions of interest (ROIs) in this study include: isocortex, hippocampus, alveus, inferior colliculus, and thalamus. The ROIs were defined based on anatomical landmarks and labeled prior to segmentation of plaque burden within the respective ROI. A machine learning pipeline to extrapolate 3D shape recognition from 2D inputs was generated by supplementing 200 desired objects (plaques) and backgrounds (non-plaque structures) respectively. Training inputs were created by using the brush tool at size 5 and 100% magnification. Two machine learning training files were generated as hemibrains with few plaques have significantly distinct plaque morphologies, thus warranting a separate training file for adequate segmentation. Next, segment colocalization was performed to delineate plaques within specific ROIs. Upon acquiring plaques as voxel objects, noise was removed by deleting objects with voxel sizes 1-10 and plane counts 1-3 as these mostly corresponded to non-specifically stained structures or residual blood as an artifact from perfusion. Lastly, object metadata and correlated features were exported as spreadsheets for further statistical analysis and representation of the data.

### Paraffin sample preparation and immunohistochemical staining

Fixed hemibrains were dehydrated in an ascending ethanol series (50%, 80%, 100%), followed by 100% isopropanol, then a mix of 50% isopropanol and 50% xylol, and finally two rounds of 100% xylol. Paraffinization was carried out through a STP 120 tissue processing machine (Leica Microsystems). Samples were then embedded in paraffin blocks on a HistoStar embedding workstation (Epredia). The paraffin-embedded blocks were then cut into 5 μm-thick coronal sections, with the resulting slices mounted onto slides and dried overnight.

Slides were first deparaffinized at 60 °C followed by incubation in 100% xylol (twice) and a 1:1 mixture of xylol and isopropanol (once) for 10 min each. Rehydration followed using decreasing ethanol concentrations (100%, 90%, 70%, 50%) and distilled water for 5 min each. Next, the slices were incubated in an antigen retrieval buffer consisting of 10 mM Tris and 1 mM EDTA (pH 9) and boiled for 10 min, while checking to keep the solution level sufficient to prevent the samples from drying out. After cooling the slices for 20 min, they were washed in distilled water for 1 min before permeabilization in 0.1% Triton X-100 in PBS for 15 min and another washing step in PBS. For Aβ labeling, an additional incubation step in a 88% formic acid solution in water for 3 min was carried out prior to two quick washes in PBS. By using cover plates (Epredia), each slide was incubated in 100 μl of 10% goat serum as the blocking solution for 1 h. Then, the samples were incubated in 100 μl of the primary antibody mixture at 4°C overnight. Primary antibodies used for immunohistochemical staining were: anti-NeuN (mouse, Millipore, 1:100); anti-RFP (rabbit, Rockland, 1:500); anti-Aβ-6E10 (mouse, BioLegend, 1:1000); anti-Iba1 (rabbit, Wako, 1:500); anti-BCAS1 (guinea pig, 445 003, Synaptic Systems, 1:250); anti-CAII (rabbit, ab124687, Abcam, 1:1000); anti-PLP-aa3 (rat, culture supernatant, 1:200), all diluted in PBS containing 10% goat serum. After washing with PBS, the samples were incubated with fluorescent secondary antibody solution for 2 h in the dark. Mixed with 10% goat serum in PBS, the following secondary antibodies were used: anti-mouse Dylight488 (donkey/goat, Thermo-Fisher, 1:1000), anti-rabbit Alexa555 (donkey/goat, Thermo-Fisher, 1:1000); anti-mouse Alexa555 (donkey/goat, Thermo-Fisher, 1:1000); anti-guinea pig Dylight650 (donkey/goat, Thermo-Fisher; 1:1000); anti-rat Alexa555 (donkey/goat, Thermo-Fisher, 1:1000). Nuclei were stained with DAPI (Thermo-Fisher, 300 nM) in PBS. Slides were washed in PBS twice for 5 min and mounted with Aqua PolyMount medium (PolySciences). Finally, slides were left to dry overnight in a 37 °C chamber then imaged through 2D fluorescence microscopy. All incubation steps were done at RT unless stated otherwise.

### Human tissue collection

Human patient samples (Control – 1 female, 2 male, age: 74±2.83 years; AD – 2 female, 2 male, age: 72.75±1.78 years) were obtained from the Neurobiobank Munich, Germany, with ethical approval from the Ethical Commitee at the Ludwig-Maximilians University in Munich, Germany. Selection of patients was performed upon Braak staging with AD patient scores ranging from Braak 5-6 and control patient scores ranging from Braak 1-3. Post-mortem interval of patients ranged between 26-51 h. APOE genotype of all control patients are 3/3, while AD patient APOE genotypes are: 3/3, 3/4, and 4/4. For ISH experiments, formalin-fixed paraffin-embedded tissue sections were used from human samples.

### *In situ* hybridization

To validate *APP* expression in human OLs and *Bace1* deletion in OLs and ExNs, we employed the RNAscope Fluorescent Multiplex assay (ACD Bio) for paraffin-embedded samples. Briefly, paraffin slices (5μm mouse and 4μm human sections) underwent deparaffinization at 60°C for 1 h, followed by incubation in 100% xylol twice for 5 min each, and 100% ethanol twice for 2 min each. This was followed by hydrogen peroxide treatment at 40°C for 15 min and a quick wash with distilled water. Slides were then boiled in the target retrieval reagent (10 min for mouse and 20 min for human sections) and carefully washed with distilled water for 15 s and 100% ethanol for 3 min. To ensure optimal hybridization, a hydrophobic barrier was created followed by protein digestion via incubation in RNAscope Protease Plus at 40°C for 15 min (mouse sections) or RNAscope Protease IV for 20 min (human sections). The slides were again washed with distilled water twice for 2 min before hybridization with the following probes: Mm-Mbp (451491-C1), Mm-Bace1-C2 (400721-C2), Mm-Slc17a7-C3 (416631-C3), Hs-MBP-C2 (411051-C2) or Hs-APP-C1 (418321-C1) at 40°C for 2 h. Slides were placed in a 5x SSC buffer for overnight storage. Signal amplification was carried out next with the provided amplification reagents followed by subsequent incubation with HRP for the specific channel probes at 40°C for 15 min. For triple visualization of mouse sections, the following fluorophores were applied: Opal 520, 570, and 690, at 40°C for 30 min. For double visualization of human sections, TSA Vivid Fluorophores (570 and 650) were used The slides were again washed and stained with DAPI (Thermo-Fisher, 300 nM) for 10 min prior to mounting with Aqua PolyMount medium (PolySciences).

Upon epifluorescence imaging, validation of *Bace1* deletion in mouse cortical ExNs was performed manually due to the presence of ample satellite OLs in the cortex. A 500 μm-wide ROI spanning all cortical layers were drawn for each coronal brain slice, more specifically in the parietal or somatosensory cortex overlying the hippocampus. For nuclei expressing *Slc17a7, Bace1* puncta was quantified to yield individual ExN *Bace1* counts. The data was grouped into distinct cortical layers which were delineated based on landmarks. The distribution of cortical ExN *Bace1* counts were plotted and the aggregated counts compared between groups. Similar manual quantification was also performed on OLs in the fimbria. For nuclei that express *Mbp, Bace1* puncta was quantified to yield individual OL *Bace1* counts. Validation of *Bace1* deletion in hippocampal ExNs was carried out semi-automatically by first drawing ROIs of the different hippocampal regions and creating a pipeline employing a nuclear detection plugin (StarDist), expanding the captured nuclear ROIs by a pixel size of 10 as not all transcripts were situated in the nuclei. Combined with particle analyzer and watershed binarization, *Bace1* puncta detected in hippocampal ExNs were also plotted with aggregated counts compared between groups.

With regards to the human data, images of the entire human brain sections were acquired with the Pannoramic Midi II Slide Scanner (3D HISTEC) with the 20x objective and smaller selected regions with the 40x objective. The human cortex was divided into three macro areas corresponding to layers 1 and 2 (L1-2), layers 3 and 4 (L3-4) and layers 5 and 6 (L5-6). For each macro area, 6-12 images of similar size were selected using CaseViewer (v2.4, 3D HISTEC) and exported in the .tif format via Slide Converter (v2.3.2, 3D HISTEC). To keep experimenters blinded for analysis, the selected images were randomized using a Fiji filename-randomizer plugin and positive cells were counted using the Fiji CellCounter plugin.

### *In vitro* OL culture and immunocytochemical staining

OPCs were isolated from p7 mouse brains using magnetic activated cell sorting and anti-NG2 MicroBeads (Miltenyi Biotec). Tissue dissection and cell sorting were performed under sterile conditions. The neural tissue dissociation kit was utilized combined according to the manufacturer’s protocol and as described^39^. In short, dissected brains were cut into smaller pieces prior to being transferred into enzyme mix 1 followed by incubation at 37°C for 15 min. Next, incubation with enzyme mix 2 was done at 37°C for 20 min with manual dissociation. Following tissue dissociation, the tubes were centrifuged at 1200 rpm for 5 min and the supernatant decanted while the pellet was resuspended in DMEM with 1% horse serum. We then passed the cell suspension through a 70 μm cell strainer to remove debris or undigested tissue prior to washing with DMEM and passing the suspension through a 40 μm strainer. The tubes were again centrifuged at 1200 rpm for 10 min and the pellet was resuspended and incubated in warm OPC culture medium consisting of 100 ml NeuroMACS media, 2 ml MACS NeuroBrew21, 1 ml Pen/Strep, and 1 ml L-GlutaMAX at 37°C for 2 h. Next, tubes were centrifuged at 1200 rpm, 4°C for 10 min followed by pellet resuspension and incubation in NG2 MicroBeads diluted in DMEM with 1% horse serum (10 μl NG2-beads per 10^7^ total cells) at 4°C for 15 min. The cell suspension was again centrifuged at 1200 rpm, 4°C for 10 min and the supernatant was removed before pellet resuspension in 5 ml DMEM with 1% horse serum. LS columns (Miltenyi Biotec) were first attached to a magnet before activating with DMEM containing 1% horse serum. The columns were washed thrice with DMEM after addition of the cell suspension. The columns were finally detached from the magnet and flushed with 5 ml DMEM containing 1% horse serum to collect bound cells. Upon detachment, samples were centrifuged at 1200 rpm for 5 min and the pellet was resuspended in proliferation medium. OPCs were plated at a density of 1.2 x10^5^ cells per well on a 12-well plate in proliferation medium, before replacement with OPC differentiation medium at DIV4. Of note, coverslips were initially placed within each well for immunocytochemical purposes. Cells were fixed at DIV8 with 4% PFA and washed with PBS thrice for 5 min each.

For immunocytochemical labeling, cells were permeabilized with cold 0.3% Triton X-100 in PBS and blocked with 10% goat serum and 0.03% Triton X-100 in PBS for 1 h. The primary antibodies anti-PLP-aa3 (rat, culture supernatant, 1:200); anti-APP-Y188 (rabbit, ab32136, 1:500); anti-BACE1-3D5 (mouse, hybridoma culture supernatant, 1:500); anti-BCAS1 (guinea pig, 445 003, Synaptic Systems, 1:250), were diluted in 1.5% horse serum in PBS and applied at 4°C overnight. Coverslips were washed with PBS thrice for 5 min and incubated in the following secondary antibodies diluted in PBS, anti-rat Dylight488 (donkey/goat, Thermo-Fisher, 1:1000), anti-mouse Alexa555 (donkey/goat, Thermo-Fisher, 1:1000); anti-rabbit Dylight650 (donkey/goat, Thermo-Fisher; 1:1000); anti-rabbit Alexa488 (donkey/goat, Thermo-Fisher, 1:1000); anti-guinea pig Dylight650 (donkey/goat, Thermo-Fisher; 1:1000), for 1 h. The samples were washed twice for 5 min before incubation with DAPI (Thermo-Fisher, 300 nM) in PBS. Lastly, cells were washed briefly in PBS prior to mounting with Aqua-PolyMount for confocal imaging. All incubation steps were done at RT unless stated otherwise.

**Epifluorescence and confocal microscopy**

For epifluorescence imaging, we used a Zeiss Observer microscope equipped with Plan-Apochromat 20×/08 and Fluar 2.5×/0.12 objectives; Colibri 5 LED light source (630 nm, 555 nm, 475 nm, and 385 nm excitation wavelengths); 96 HE BFP, 90 HE DAPI, GFP, Cy3, 38 GFP, 43 DsRed, 50 Cy5 Zeiss filter sets; Axiocam MrM and SMC900 motorized stage. In the ZEN imaging software (ZEN Blue v3.3, Zeiss), a preview scan was first taken at 2.5× magnification to draw the ROI outline and focus on the brain slice. Upon switching to 20× magnification, support points were distributed using the onion skin method and manually focused in a pre-selected reference channel. Acquired via the single channel mode, the resulting tiled images were stitched in ZEN and pseudocolours (white, blue, and yellow) were assigned to different channels.

For confocal microscopy, images were partially acquired via the Zen software with a Zeiss LSM 800 Airyscan confocal microscope equipped with Plan-Apochromat 63x/1.4 oil DIC M27 objective. Alternatively, images were acquired via the LasAF software with a Leica SP8 Lightning confocal microscope equipped with an argon laser and a tuneable white-light laser with 63x/1.4 glycerin objective. Both confocal microscopes are situated at the European Neuroscience Institute and MPI-NAT, City Campus respectively.

### Analysis of 2D microscopy images

All 2D image analysis was performed on Fiji (v.1.53c)^40^. For validation of CNP-Cre specificity in the cortex and hippocampus, thresholding and particle analyzer were performed to segment and quantify neurons, OLs, and RFP+ cells. Quantification of RFP+ OLs in the CC, however, was performed manually due to the dense amount of OLs in WM tracts. Quantification of 2D Aβ and microgliosis first started with segmentation of ROIs of the brain prior to thresholding and measurement of positive area. Microscopic analysis of OL numbers between controls and cKOs similarly started with brain ROI segmentation followed by thresholding and particle analyzer. As for OL numbers in WM tracts, manual quantification was again performed. Finally, percentage ROI area of the cortex and hippocampus occupied by myelinated structures were obtained upon thresholding and mean intensity values of major WM tracts were measured. All statistical analyses are described in more detail in the respective figure legends with all relevant p-values shown within each graphical representation.

### Electron microscopy

To prepare samples for electron microscopy, fixed brains were first immersed in ice-cold PBS and cut sagittally into 300 μm-thick slices with a vibratome (Leica VT1200). The caudal corpus callosa alongside adjacent tissue were punched with a 2 mm diameter punching tool and embedded in Epon (EM TP, Leica) prior to conventional embedding steps. Samples were first incubated at 4°C in 0.1 M phosphate buffer thrice for 15 min each, 2% OsO_4_ for 4 h, and 0.1 M phosphate buffer thrice for 10 min each. The subsequent steps involved treating samples with an ascending acetone bath concentration of 30%, 50%, 70%, 90% for 20 min each and four changes of 100% acetone for 10 min each. The samples were incubated in increasing Epon concentration in acetone, first 33% and 50% for 2 h and lastly in 66% and 100% Epon for 4 h. Upon completion, samples were covered with an Epon-filled gelatin capsule prior to polymerization at 60°C for 24 h. Epon-embedded samples were trimmed and cut to produce semi-thin 500 nm sections on Leica UC7 ultramicrotome (Leica, Vienna, Austria) equipped with a diamond knife (Diatome). Semi-thin sections were stained with Azure II for 1 min to stain lipid-rich areas to confirm the region of interest. Next, 60 nm-thin sections were cut and transferred to Formvar-coated copper mesh grids (Science Services) before air drying. Uranyless contrast was applied onto the sections for 30 min (Electron Microscopy Sciences) and washed five times with distilled water. Samples were air dried before further imaging. At least 10 digital pictures were captured at x4000 magnification with the Zeiss EM900 for ultrastructural analysis.

Electron micrographs of the caudal corpus callosa were analyzed with Fiji. We first quantified the number of myelinated axons from at least five micrographs per animal with the cell counter plugin. These amounts were then averaged and used for comparison between groups. To conduct g-ratio analysis, we measured the axonal and myelinated fiber areas of at least 100 fibers per animal. A 5 × 5 grid was applied to each image and only myelinated fibers crossing the grids were quantified to avoid bias. The axonal area was then divided by the whole myelinated fiber area to generate the g-ratio, and this value was plotted against the axon caliber to observe linear regressions and data spread between groups.

### Protein fractionation

Preparation of insoluble and soluble fractions from mouse brain tissue was carried out based on a previously described protocol^29^. Briefly, a tissue homogenizer (Precellys) was utilized to homogenize microdissected cortical and CC fractions in reaction tubes containing ceramic beads in pH 8.0 cold lysis buffer (700 μl for cortex and 500 μl for CC) containing 120 mM NaCl, 50 mM Tris, cOmpleteTM protease inhibitor cocktail, and phosphatase inhibitor cocktail 3 (1:100 dilution). The following settings were used for the homogenization at 4°C: 6500 g twice for 30 s with a 15 s pause in-between. The homogenate was carefully transferred to a 1.5 ml reaction tube prior to spinning with a bench-top centrifuge (Eppendorf) at 17000 g, 4°C for 20 min. The supernatant was collected and served as the soluble protein fraction while the pellet was resuspended in 2% SDS (500 μl for cortex and 300 μl for CC). The solution was then sonicated on ice for 1 min until the pellet completely dissolved. To remove DNA, 1 μl of benzonase was added into the solution and incubated at RT for 5 min. The samples were again centrifuged at 17000 g, 4°C for 20 min before transferring the supernatant to a fresh collection tube, serving as the insoluble fraction. Both fractions were stored in -80°C for further use.

### Western blotting

Only the insoluble fraction was utilized to probe for APP processing machinery via Western blotting. Protein concentration of each lysate was determined via detergent-compatible protein assays (Bio-Rad). Samples were then mixed with a commercial 2x Tris-tricine sample buffer, 0.05 M DTT, and an equal amount of protein to create an array of probes with equal protein concentration (2 μg/μl for cortex and 1 μg/μl for CC). Typically, 25-30 μl of working lysate was loaded per lane in precast 10-20% Tris-tricine SDS–PAGE gradient gels (Novex). Electrophoresis was performed at 120 V for 2 h. Protein transfer was done using the Bio-Rad wet-blotting system at 500 mA for 1 h on 100% methanol-activated low-fluorescent Immobilon-FL membranes (0.45 μm pore size, Merck). The blot transfer buffer consisted of 25 mM Tris, 190 mM glycine, and 20% methanol. After a quick wash with distilled water, total protein amount for normalization was measured via a Fastgreen stain on the blot. Washed membranes were transferred to 0.0005% Fastgreen FCF (Serva) in a washing solution containing 30% methanol and 7% glacial acetic acid in distilled water for 5 min. The membranes were then washed twice for 1 min each in the washing solution prior to visualization using a ChemoStar fluorescent imager (Intas). Upon completion, membranes were washed thoroughly in TBS with Tween (TBS-T, 0.05%) at least thrice for 10 min each. We then blocked the membranes in 5% BSA in TBS-T at RT for 1 h before an overnight incubation in primary antibody solutions containing 5% BSA in TBS-T at 4°C on a rotating shaker. The following primary antibodies were used: anti-C-terminal APP (rabbit, 127-003, Synaptic Systems, 1:500); anti-BACE1-3D5 (mouse, hybridoma culture supernatant, 1:500); anti-PSEN1 (rat, MAB1563, Sigma-Aldrich, 1:500). The next day, membranes were washed in TBS-T thrice for 10 min each prior to incubation in secondary antibody solutions also containing 5% BSA in TBS-T. The following secondary antibodies were used: anti-rabbit IgG (H+L) IRDye 800 (Licor, 1:5000); anti-mouse IgG (H+L) IRDye 680 (Licor, 1:5000); anti-rat Dylight650 (donkey/goat, Thermo-Fisher; 1:5000). Upon washing in TBS-T thrice for 10 min each, membranes were scanned using an Odyssey platform (Licor). For quantification performed on raw images, background was subtracted and bands were analyzed using Fiji (integrated density). Target protein content was normalized to the Fastgreen bands respective controls as indicated in the graphical representations, alongside the statistical analysis performed.

### Electrochemiluminescence Aβ immunoassay

To determine Aβ levels in specific brain regions, we turned to the V-PLEX Plus Aβ Peptide Panel 1 (6E10) kit (Meso Scale Discovery) and conducted experiments based on instructions provided by the manufacturer. The kit allows multiplex measurement of Aβ38, 40, and 42 from single wells. First, 150 μl of Diluent 35 was added into each well for blocking before the plates were sealed and incubated with shaking at RT for 1 h. Each well was subsequently washed thrice with 150 μl of wash buffer containing 0.05% Tween 20 in PBS (PBS-T). From a detection antibody solution containing 50x SULFO-TAG Anti-Aβ 6E10 antibody diluted in Diluent 100, 25 μl was added into each well followed by addition of 25 μl samples or calibrators per well. The plate was again sealed and incubated with shaking at RT for 2 h. Each well was again washed thrice with 150 μl PBS-T (0.05% Tween 20 in PBS) before addition of 150 μl of 2x Read Buffer T. Lastly, plate measurement was carried out via the MSD QuickPlex SQ 120 reader. In all assays performed, two technical replicates of samples and calibrators were included.

#### Acknowledgments

The authors would like to thank all personnel of the animal facility of the Max Planck Institute for Multidisciplinary Sciences (MPI-NAT), City Campus; A. Fahrenholz for her technical contribution in sample preparation for biochemical and imaging experiments; B. Sadowski for sample preparation for electron microscopy; members of the KAGS subgroup and the Department of Neurogenetics for scientific input; and M. Thalmann for his inputs on statistical analysis and data segmentation. Work pertaining to this manuscript was performed in the laboratory of K.-A.N. and supported by the Deutsche Forschungsgemeinschaft (DFG, TRR274), the Dr Myriam and Sheldon Adelson Medical Foundation (AMRF) and an ERC Advanced Grant (MyeliNANO). We extend our gratitude to the Gene Expression Omnibus (GEO) and AD Knowledge Portal data platforms. The valuable data provided by these platforms owes its existence to the dedication of research volunteers and the collaborative efforts of contributing researchers.

## Author contributions

A.O.S., C.D., and K.-A.N. conceptualized and designed the experiments. A.O.S. and T.N. planned and performed imaging and biochemical experiments. T.S. arranged and analyzed snRNA-Seq datasets. X.Y., S. Siems, and S.E. executed *in vitro* culture experiments. C.B. prepared samples for imaging and executed biochemical experiments. E.C.O. carried out biochemical experiments. Z.W., T.R., and W.M. prepared samples and advised on electron microscopy experiments. B.B. assisted in developing an automated analysis pipeline for ISH quantification. B.B. and S. Siems performed confocal imaging. B.M.M. guided and assisted with electrochemiluminescence assay. S. Su amanian assisted in data analysis. F.B. and K.O. performed all genotyping. L.E. and S.J. performed and analyzed ISH on human tissue. L.S. and S.A.B. conducted pilot biochemical experiments. K.S. and R.V. supplied an antibody for imaging and biochemical experiments. S.G. provided *Nex-Cre* mice. T. Saito and T. Saido provided *APP*^*NLGF*^ mice. R.Y. provided *Bace1*^*fl/fl*^ mice. G.C.-B. accommodated infrastructure for computation of some snRNA-Seq datasets. H.-W.K. and O.W. advised on electrochemiluminescence assay. S.J. provided and advised on human ISH data. A.O.S. analyzed data and constructed figures. A.O.S., C.D., and K.-A.N. prepared the manuscript.

## Competing interests

The authors declare no competing interests.

## Notes

https://github.com/TSun-tech/2023_Sasmita_OL_BACE

### Competing Interest Statement

The authors have declared no competing interest.

